# Widespread, perception-related information in the human brain scales with levels of consciousness

**DOI:** 10.1101/2022.09.19.508437

**Authors:** Andrew D. Vigotsky, Rami Jabakhanji, Paulo Branco, Gian Domenico Iannetti, Marwan N. Baliki, A. Vania Apkarian

## Abstract

How does the human brain generate coherent, subjective perceptions—transforming yellow and oblong visual sensory information into the perception of an edible banana ^1^? This is a hard problem. The standard viewpoint posits that anatomical and functional networks integrate local, specialized processing across the brain to somehow construct unique percepts. Here, we provide evidence for a novel organizational concept by uncovering task-specific information distributed across the human brain. First, we show that functional magnetic resonance imaging (fMRI) can uncover task-specific information throughout the neocortex, even across voxels traditionally discarded as “noise” (t-statistics ≈ 0), challenging the sensitivity of traditional linear, univariate analytical approaches. Remarkably, task-specific signals could also be uncovered from across-subject variances and were ubiquitous even in the subcortex and cerebellum. Finally, we show that the widespread signal in regions remote from a task’s primary and secondary sensory cortices depends on the level of sedation, suggesting it is related to perception^†^ rather than sensory stimulus encoding. We hypothesize that these widespread, task-specific, and consciousness level-dependent signals may be the basis for coherent, subjective perceptions.

## MAIN TEXT

fMRI has transformed how we study the brain, allowing the non-invasive measurement of correlates of neural activity on the scale of millimeters. This high spatial resolution enables the comparison of blood oxygenation level-dependent (BOLD) activity within and between tasks to unravel the functional properties of local neural circuits. Such task-based fMRI studies commonly use forward inference to identify task-related brain areas: they rely on the correlation between each voxel’s activation time-course with the task’s temporal profile. Subsequently, contrasting brain activity maps of different tasks generates a contrast map, which yields task-specific localized blobs when thresholded. The standard assumption of these mass-univariate analyses is that only the identified blobs reliably contain task-specific information ^2^.^‡^ In this report, we challenge this assumption by showing the reliable presence of task-specific information throughout the entire neocortex—including regions identified as “noise” by univariate measures (*t*-statistics ≈ 0)— which is uncovered by integrating over large swaths of voxels (~10,000 voxels). After discovering pan-neocortical information content, we probe for and uncover task-specific information in its across-subject variance, and in addition, in the subcortex and cerebellum. Next, we assess how different levels of sedation perturb the presence of information. We show that the omnipresent information degrades with increasing levels of sedation. Rather than being task-specific, this brain-wide spread of information appears to reflect perceptual (conscious) processes and may be involved in extracting subjective, wholistic concepts from incoming sensory inputs, as in identifying the edible banana.

Decoding can be used to assess the information contained in neuroimages by transmuting brain activity into a single number that is, ideally, monotonically related to a task of interest. This monotonicity facilitates the discrimination between the tasks of interest and no interest, the performance of which indicates the amount of task-specific information in the data. Based on six datasets with different sensory stimuli (Table S1; *N*=293 subjects) ^3–8^, we built simple decoding models using only the *t*-statistics from mass-univariate contrasts (Fig 1, top left; Fig S1). The first part of our study examines decoding for five of these datasets, four of which contain two stimuli and one of which includes four stimuli, totaling ten different stimulus pairs or contrasts 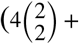 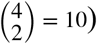. We binned the *t*-statistics by magnitude to create ten decoders for each contrast; the first decoder comprised of voxels with the highest *t*-statistics (10^th^ decile) and the last decoder comprised of voxels with the lowest *t*-statistics (1^st^ decile) (Fig 1, top right). We tested each decoder by calculating the dot product between the decoder (sign, magnitude, and location of *t*-statistics within a single decile; e.g., Fig 1, bottom right) and each brain activity map (general linear model (GLM)-derived maps of parameter estimates), yielding a weighted sum of task-related activity across all voxels comprising the decoder (Fig 1, bottom middle). We used the 0.623+ bootstrap to obtain unbiased estimates of between-subject areas under the receiver operating characteristic curve (AUC) as an indicator of discrimination performance (Fig 1, bottom left). To succinctly describe our results, we meta-analyzed the resulting AUCs and their bootstrapped variance-covariances (*Methods*).

**Figure 1.**
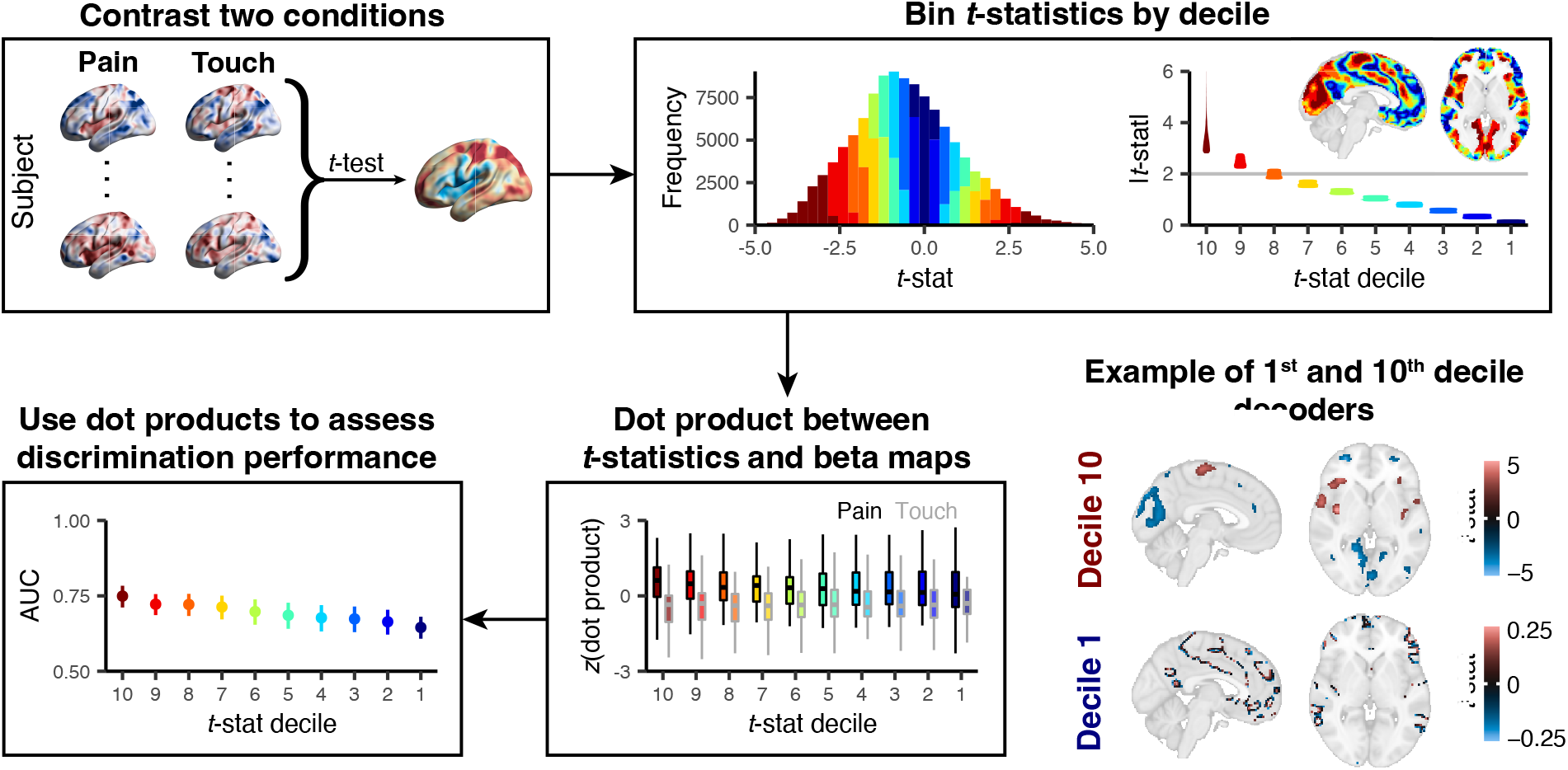
Generation of decoders to assess the presence of task-related signals. Voxel-wise paired *t*-tests were performed on subjects’ brain activity maps using a task of interest (e.g., pain) and no interest (e.g., touch). These *t*-statistics were then binned into deciles based on their absolute magnitudes. The grey line depicts the standard minimum threshold used to dichotomize brain regions that discriminate between tasks (univariate “statistical significance” at *α* = 0.05; uncorrected for multiple comparisons). Each decile of *t*-statistics and their locations in the brain became a decoder. On the bottom right, we show examples of these decoders for the 10^th^ (comprised of large blobs) and 1^st^ deciles (scattered voxels). Although the deciles were derived using the absolute value of *t*-statistics, the decoders incorporated the *t*-statistics’ signs. We then calculated dot products between the decoder derived from each decile and brain activity maps. These dot products are analogous to ‘linear predictors’ from a regression model. Here, we *z*-scored the dot products within each decile for visualization purposes. We calculated AUCs based on these dot products, where higher dot products were assumed to correspond to the task of interest. We used the 0.632+ bootstrap to obtain unbiased AUCs when testing our decoders.

### Task-specific information is widespread across the human brain

Decoding performance was consistently above chance (AUC > 0.5) for all deciles across nine out of ten contrasts. Despite univariate *t*-statistics in the lowest decile being close to zero, decoding performance was only marginally poorer in the lowest decile than in the highest decile (Fig 2, Table S2). Therefore, regions in neocortical grey matter commonly thought to be orthogonal to the task in univariate analyses contain robust task-related information. These findings complement recent work using statistical learning to optimize voxel weights for predictive performance ^9–13^ and demonstrate that the presence of information is far more distributed across the brain than previously thought. Moreover, our analyses establish for the first time how accessible this information truly is: our models use mass-univariate *t*-statistics without any regularization or consideration of the *t*-statistics’ joint distribution. Regularization and multivariable modeling are not necessary, and even voxels with *t*-statistics close to zero can jointly discriminate tasks from one another quite well (meta-analytic AUC > 0.7). Therefore, our results indicate the presence of task-related information throughout the neocortex, which degrades slowly as a function of the univariate signal-to-noise metric (*t*-statistic deciles).

**Figure 2.**
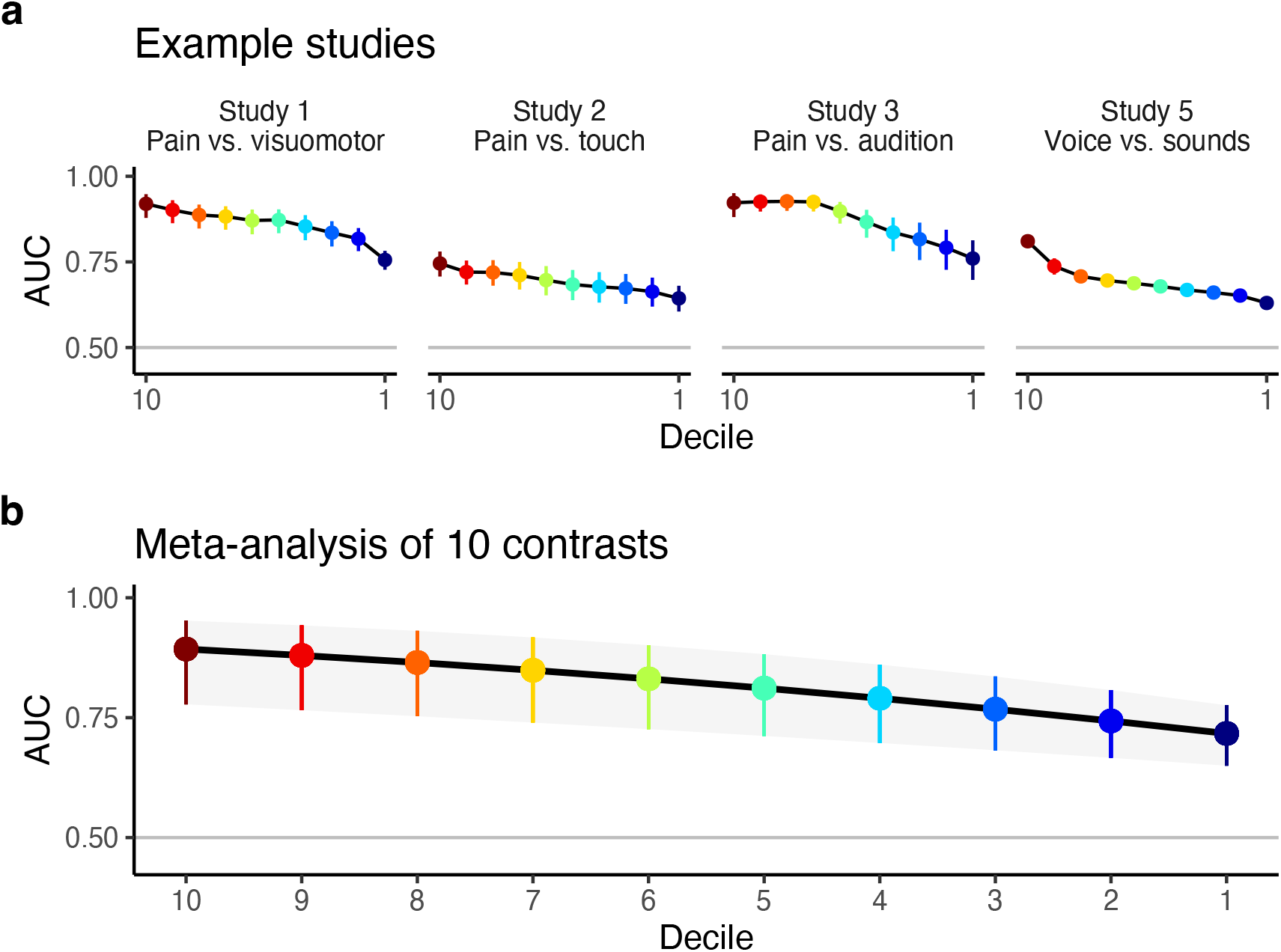
Voxels across the entire neocortex contain task-specific information. (**A**) Four example studies of neocortical decoding performance across all *t*-statistic deciles. Although all four studies have task-specific information in all 10 deciles, the degree to which the tasks can be discriminated differs slightly between studies. Error bars indicate ± SE. (**B**) A mixed-effects meta-analysis across 10 contrasts reveals that all ten deciles can, on average, discriminate between tasks. The ability to discriminate between tasks implies the presence of task-specific information, meaning that even voxels with *t*-statistics close to zero (decile 1) contain marked task-specific information. Error bars indicate ± CI_95%_.

When decoding tasks from neuroimages, one cannot draw inferences about the causal role of the predictors (voxels) in the decoding outcome (task) ^14^. There are many reasons why information may be present in acausal structures. For example, physiological and non-physiological noise may have task specificity ^15^. We attempted to rule out such confounds by decoding tasks using white matter and cerebrospinal fluid (cf. grey matter). In general, decoding performance was poor in these regions (Fig S2), suggesting these negative controls contain less signal than the neocortex. In addition, white matter and cerebrospinal fluid decoding performance substantially covaried (*r* = 0.7), but they only weakly covaried with grey matter, implying vastly different signal sources between these structures (Fig S2). Finally, head motion artifact—another potential candidate of task-related noise that contaminates the BOLD signal—only minimally correlated with decoder responses; orthogonalizing the decoders’ dot products to head motion only slightly decreased discrimination (maximum ΔAUC < 0.05). Thus, our observed effects are unlikely attributable to task-specific, non-neural confounds.

To assess the general sensitivity of the decoders, we built them using different numbers of voxels and different amounts of added noise. Decoders with fewer voxels performed poorly and were more sensitive to added noise (Fig S3). As one might expect, the ability to successfully decode using voxels with low *t*-statistics is principally attributable to the number of included voxels. There is a continuum of explanations for why this might be the case. On one extreme, each voxel may contain a tiny amount of task-specific information. Integrating many small bits of information allows one to accumulate enough information to discriminate between tasks. On the other extreme, since *t*-statistics are empirically derived and thus will not be stable across samples, “signal” voxels may mix with “noise” voxels, creating instability in our deciles. In this case, by sampling more voxels, we are more likely to capture “signal” voxels in our “noise” decile(s) to enable successful decoding.

Our *t*-statistic analysis relies on the task’s so-called main effect within each voxel. However, task-relevant information may exist beyond the main effect—namely, in the variance structure. We evaluated the presence of higher-order, task-specific information by performing principal components analysis (PCA) on data from one of the larger studies with equisalient stimuli (Study 2, pain vs. touch; *n* = 51) 5. We extracted four principal components (PC) (Fig 3a,b), with which we decoded across-subject variance. We fit PCs in the training sample to assess the across-subject variance-covariance structure of the pain condition. Next, we projected the test sample’s pain and touch activation maps into the same PC space (Fig 3c). By comparing the multidimensional structure of the test samples to the original training sample, we could distinguish the pain activation maps from the touch activation maps 93% of the time. Interestingly, this suggests unique patterns of task-related activity (PCs) differ across participants. From a constructivist viewpoint, one interpretation is that there is a many-to-one relationship between brain activity and gross percepts, consistent with Lisa Feldman-Barrett’s notion of *degeneracy* ^16^. In our view, a related idea from motor control, *abundance*, expands on degeneracy by providing a teleological explanation: since a one-to-one relationship between neural activity and percepts would be overly prescriptive and, as a result, inflexible and unstable, a many-to-one relationship allows for the nervous system to organize and tune itself as needed to complete a task ^17^—in this case, generating a percept. Many different combinations of brain activity patterns may be sufficient to create the perception of a banana. Since these discordant patterns are generalizable, individual difference research may be capable of discovering neurocognitive rules. This analysis is independent of the *t*-statistic decoding (Fig 3d) and demonstrates the presence of ample task-specific information in the higher moments of brain activity maps.

**Figure 3.**
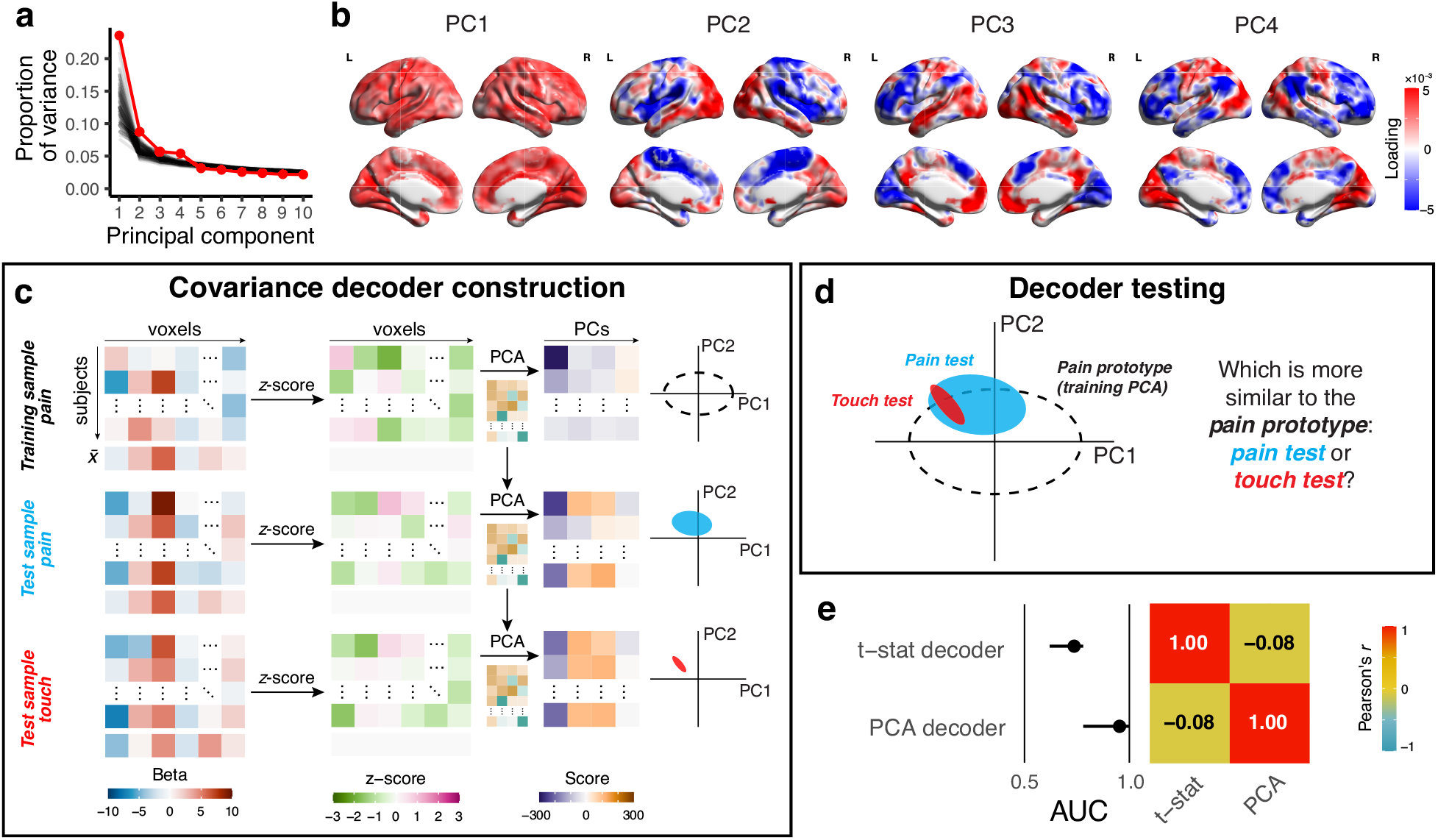
Principal components analysis reveals task-related covariance across participants that is orthogonal to the t-statistic decoding. (**A**) We performed principal components analysis (PCA) across subjects using the brain activity maps for the pain condition (Study 2).^5^ Parallel analysis indicated that the first four principal components (PCs) were above the noise floor. (**B**) Each PC has a unique corresponding spatial structure; their covariance structure is neither strictly local nor contiguous. (**C**) We used the PCs from the pain condition as a decoder (a “prototype”) by *z*-scoring the brain activity maps from out-of-sample pain and touch conditions. Next, we projected them onto the four PCs from the training sample. 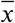 indicates the column mean, which is transformed to zero when the brain activity maps are *z*-scored; *i.e*., all data that go into the PCA are centered and thus have a mean of 0. (**D**) We compared the out-of-sample PC score distributions to the pain prototype distribution using Kullback-Liebler divergence. Whichever test sample had the lower divergence “won”, and the relative frequency of wins in the pain condition was the forced choice AUC. (**E**) The PCA decoder outperformed the *t*-statistic decoder (AUC = 0.95 vs. 0.74). This favorable performance of PCA is partly attributable to the use of the entire sample to calculate a forced choice AUC while the *t*-statistic uses individual participant scores to calculate a continuous AUC. Importantly, the AUCs were approximately orthogonal (*r* = −0.08) indicating that they decoded using unique sources of information.

Next, we tested whether the *t*-statistic information content is specific to the neocortex. Repeating our analyses in the subcortex and cerebellum revealed that information is present throughout both regions, even where *t*-statistics are approximately zero (Fig S4, Fig S5). Cerebellar information varied more between task pairs than the neocortex (three examples shown in Fig 4A). These results complement recent work by Nakai and Nishimoto ^18^, who used the subcortex and cerebellum to decode 103 cognitive tasks using a within-subject approach based on more complex models trained using statistical learning. In contrast, we used *t*-statistics from regional activity maps to decode across-rather than within-subjects. Our meta-analysis across task contrasts showed that the performance of the subcortical and cerebellar decoders was only slightly inferior to that of the neocortex-based decoders, even after controlling for the number of voxels (Fig 4B, Fig S6). Overall, we observe that subcortical and cerebellar structures contain widespread, task-specific information, evidencing that information spread is not restricted to the neocortex but is present across the entire human brain.

**Figure 4.**
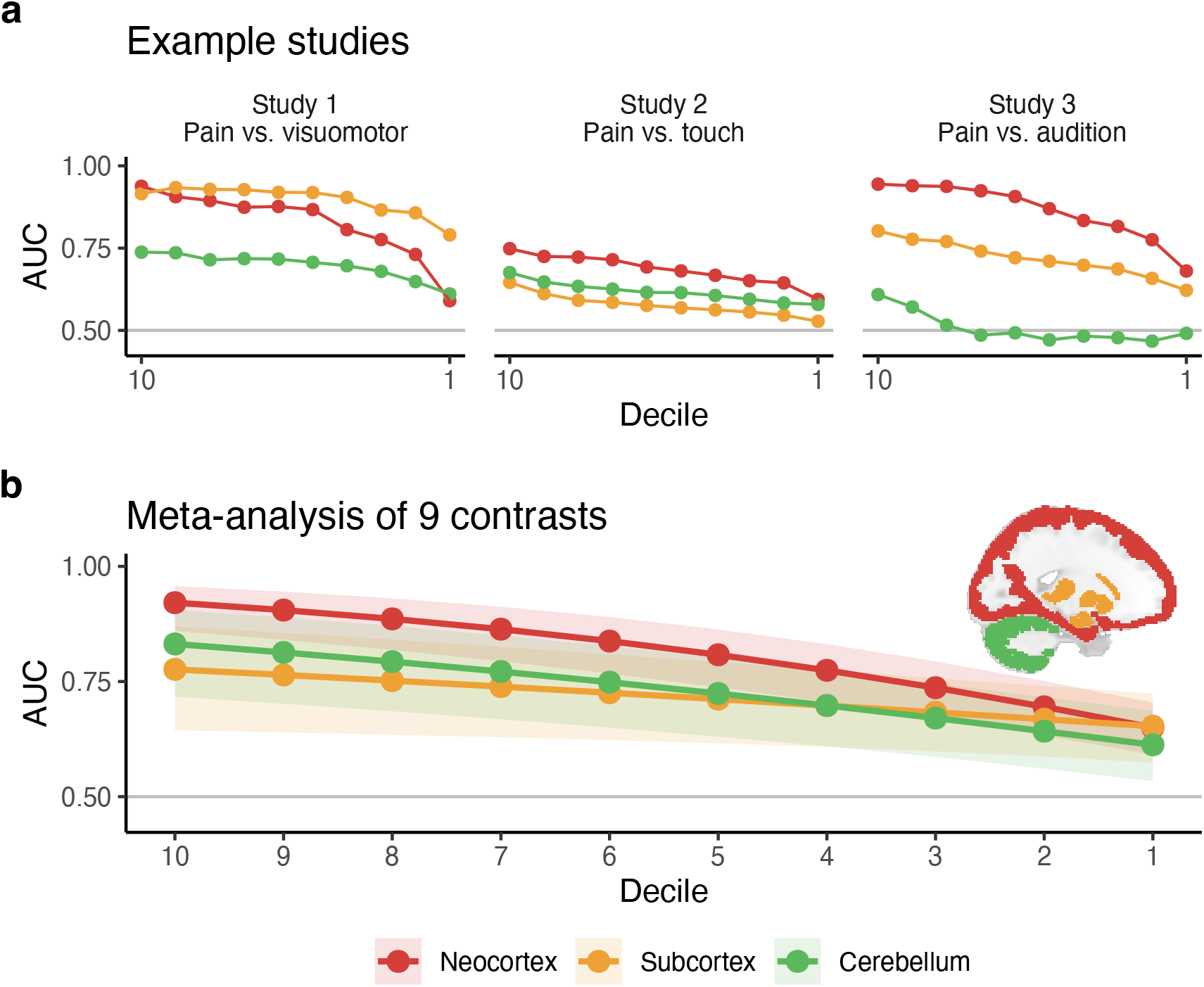
Task-relevant information is pervasively present throughout the subcortex and cerebellum but to a lesser extent than in the neocortex. (**A**) Three example studies demonstrate marked differences in regional task-specific information. In Study 1, the relative task-specific information in the neocortex shifts from being closer to the subcortex to the cerebellum. In Study 2, all three regions are similar, while in Study 3, there is a consistent pattern with the three areas being starkly different. (**B**) After controlling for the number of voxels in the neocortex, subcortex, and cerebellum, a meta-analysis across 9 contrasts (excluding Study 5) reveals that task-related information exists across all regions and deciles. Task-related information in the neocortex dominates for higher deciles, but this superiority vanishes in smaller deciles. Error ribbons indicate ± CI_95%_.

The notion of widespread, task-specific cortical dynamics is gaining traction across multiple fields of neuroscience. Human fMRI work demonstrates that 100 repetitions of the same task (three participants, 9–10 sessions over three months) can uncover neocortex-wide information ^19^. Decoding studies, which rely on statistical learning approaches, evidence the existence of task-specific information outside of GLM areas ^9–13^; it has also been demonstrated that some of these local information patterns can be uncovered via multivariate decoding methods ^20–22^. Similarly, recent human and macaque monkey fMRI studies demonstrate the presence of retinotopic tuning in cortical and, in macaques, subcortical regions remote from the visual cortex ^23,24^. To complement fMRI evidence, wide-field calcium imaging and Neuropixels recordings in rodent models capture mesoscopic neocortical and subcortical dynamics on a moment-by-moment basis, revealing brain-wide, task-specific activity across several cognitive domains ^25–28^. Our results complement this prior work by demonstrating the presence of brain-wide, task-specific information in human brain fMRI and the ease with which this information can be extracted from the mean and variance of brain activity.

Our findings have profound implications for task-based fMRI analysis. Null hypothesis significance testing (NHST) is the dominant statistical paradigm in task fMRI studies, which involves drawing dichotomous inferences from mass univariate GLM analyses: Is a voxel “activated” or not? Sub-threshold voxels are discarded despite that many of them contain task-specific information. Thus, NHST masks task-related activation to maintain type I error rates (*e.g*., *α* = 0.05) ^19^. Moreover, since fMRI meta-analyses typically rely on “vote-counting” procedures ^29,30^, regions with small, consistently subthreshold effects will not be uncovered. However, these inferential issues can be at least partially overcome. On the study level, dichotomous interpretations of results, including those intrinsic to NHST, should be avoided. Analytically, taking advantage of covariance in the data and more flexible functional forms (cf. linear effects), such as the inclusion of basis or nonparametric functions, nonlinear terms, and/or temporal derivatives, may also improve sensitivity ^31–33^. On the meta-analytic level, data sharing can facilitate mega-analyses, enabling researchers to pool raw data from many studies, and sharing unthresholded maps of the estimated effects and their standard errors can facilitate proper meta-analysis (cf. vote-counting) ^29^. Evidently, there is much information left on the table in task fMRI studies.

Understanding the nature of this brain-wide information is more challenging than identifying its existence. Recent work in mice demonstrates widespread cortical dynamics to be necessary for behavior—preventing local clusters of activation impairs performance, suggesting a functional rather than epiphenomenal role ^26^. If activation across the entire brain is necessary for task performance, it is more likely that the information we detected across brain regions is complementary than redundant. In other words, different brain regions capture distinct properties of the task. However, we remain agnostic regarding the role of this widespread information in conscious perception instead of simply being a task correlate. To address this, we will now link these findings to consciousness.

### Widespread, task-specific information scales with consciousness

Neurophysiological theories of consciousness rely on brain-wide information sharing, which is posited to be necessary but not sufficient for consciousness ^34^. Conceivably, the association between information sharing and consciousness ^35^ suggests that task-specific brain-wide information should attenuate with increasing levels of sedation. Information cannot be omnipresent if it is not readily shared across the brain. But how do states of consciousness interact with task-specific, brain-wide information content? To assess this, we analyzed a dataset in which individuals listened to an auditory stimulus (five-minute audio from a movie) under different levels of sedation ^7,8^. Since there was no auditory task vector, we averaged participants’ auditory cortex time courses to serve as the task vector. We used a separate resting-state scan as a negative control. Consistent with our analyses above, *t*-statistic decoding showed that task-related information was omnipresent across the neocortex when participants were awake. However, this information degraded with increasing levels of sedation and was partially restored while recovering from sedation (Fig 5A and B). We performed a region-of-interest (ROI)-based analysis to complement the decile analysis. In the awake state, different regions exhibited distinct abilities to discriminate the task from resting-state, with the auditory cortex exhibiting the greatest discrimination. Moreover, the auditory cortex’s task-specific signal was invariant to sedation level, but task-specific information degraded with deeper levels of sedation across all other ROIs (posterior, anterior, visual, and motor cortices) (Fig 5C). These results imply that brain-wide, task-specific information content is related to the perception rather than the encoding of the sensory stimulus.

**Figure 5:**
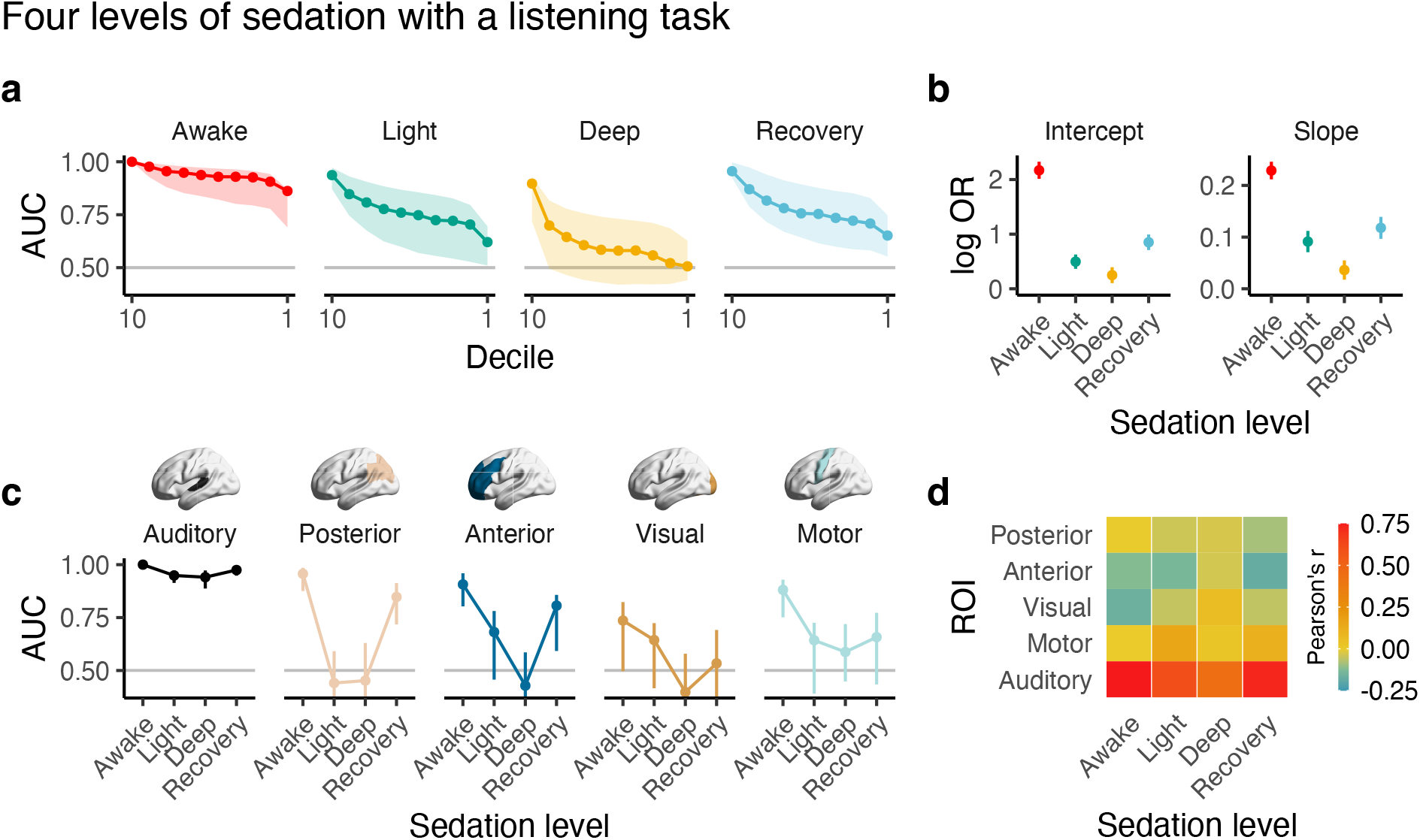
Sedation affects the neocortical distribution of task-relevant information in a region-specific manner. (**A**) Widespread cortical task-relevant information decreases with deeper levels of sedation, as indicated by the decrease in AUCs from awake to light to deep and the increase in AUC from deep to recovery. (**B**) Intercepts (defined by decile = 1) and slopes of the curves in (**A**) reveal stark decoding differences between levels of anesthesia. Awake’s first decile (intercept) has the strongest performance of the different levels of sedation. Its slope (on the logit scale; OR = odds ratio) is also greatest since it is more difficult to improve performance as AUC → 1. (**C**) The primary and secondary sensory cortices (auditory cortex) retain information across sedation levels. In contrast, other cortical regions’ information content drops off with increasing levels of sedation. (**D**) Task-specific intersubject functional connectivity decreases with deeper sedation. All error bars indicate ± CI_95%_.

The brain’s modularity and interconnected functional and structural networks must underly our observed brain-wide distribution of information ^36–39^. In particular, long-range connections and small-world networks to which they give rise provide a mechanism for efficient information sharing. The properties of these networks are thus likely to be critical for how information is communicated and captured across the brain. We elucidated the relevance of functional connectivity to our findings by examining inter-subject functional connectivity (ISFC) using the auditory cortex as a seed. We quantified the temporal relationship between a participant’s auditory cortex and all other participants’, say, posterior cortex. ISFC decreased with more sedation, much like decoding performance (Fig 5D). However, within-subject functional connectivity remained largely unperturbed (Fig S7). Thus, ISFC seems to track sedation-dependent perceptual states.

Our results complement previous work in which transcranial magnetic stimulation (TMS) was used to induce electrical potentials that differentially propagated over the neocortex as a function of consciousness state ^35^. Here, we leveraged passive sensory stimuli, but our findings are consistent: loss of consciousness degrades brain-wide information content via a downregulation in corticocortical information sharing. Therefore, consciousness seems a necessary condition for the presence of widespread task-related cortical information. Practically, our simple decoding approach may be sufficient to identify neural correlates of consciousness using natural sensory stimuli without artificially stimulating the brain (cf. TMS ^35^).

Task-related confounds, such as head motion, are likely greater when individuals are awake. However, our sedation level-dependent findings were unrelated to head motion (Fig S8). Moreover, we observed similar results whether we used auditory cortex activity from the awake or deep anesthesia conditions (Fig S9). This latter point is remarkable: auditory cortex activity with deep sedation is arguably “purer” than that in the awake condition, as higher-level processes and feedback loops will not modulate it, and similarly, head motion should be negligible. The generalizability of our findings across participants and levels of sedation reinforces that our findings represent consciousness-related neural information rather than task-related artifacts.

The neocortex is not the only neural structure involved in consciousness. Much has been discussed and debated regarding the role of the thalamus, other subcortical structures, and the cerebellum ^34^. Like the neocortex, the subcortex’s task-specific information content demonstrated a dose dependence on the level of sedation. Similarly, so did the cerebellum, although its dependence on sedation level displayed a more complex relationship (Fig S10). We should emphasize that the presence of brain-wide information reflects the state of consciousness rather than demonstrating its mechanism(s).

### Concluding Remarks

Our results render the presence of brain-wide information indisputable. We decoded this information using both the magnitude (ignoring individual differences) and variance (based entirely on individual differences) of brain activity. In doing so, we elucidated the ease with which this information can be uncovered, even in brain regions that mass-univariate analyses suggest are approximately orthogonal to the tasks being studied. We also show that the ubiquity of this information is not without bounds—consciousness is itself a necessary condition for the brain-wide spread of task-related information. Many tasks we compared are trivially simple, only involving passive sensory stimuli; yet, related information was spread across the entire brain. Neurocognitively, our results imply that perceptual states engage the entire brain. We speculate that the details of the distribution of information may define the nuanced properties of perception; for example, the edibility of the oblong, yellow object. Finally, these results strongly challenge the notion of localization of information in the brain without precluding regional specialization of function. For example, although language-specific information can be uncovered across the entire neocortex ^40^, the functional role of Broca’s area is incontrovertible ^41^. Unraveling the unique contribution of diverse brain regions to perceptual states requires future investigation, including the necessity of some and the sufficiency of others, and requiring methods beyond traditional linear, univariate analyses.

## MATERIALS AND METHODS

### Datasets

Six datasets were used in this paper; all are part of published studies and were either provided by their authors (Datasets 1–4) or downloaded from public repositories (Datasets 5 & 6). Datasets 1–4 consist of voxel-wise, whole brain, task-dependent general linear model (GLM) analysis activation maps (ftp://openpain.org/LimitsDecoding/Data/Beta_maps/). Datasets 5 & 6 consist of BOLD timeseries which were processed using standard fMRI pre- and post-processing methods described below.

#### Dataset 1

Fifteen (15), right-handed, adult subjects (mean age: 35 ± 11 years, 7 females). Subjects had no history of pain, psychiatric, or neurological disorders. fMRI data were collected while subjects received thermal stimuli across 3 temperatures: 47, 49, and 51°C. Subjects continuously rated, using a finger span device ^42,43^, their pain from 0 (not painful) to 100 (worst imaginable pain) (“pain rating” task). A control scan was performed while subjects used the finger span device to track a moving bar projected on the screen (“visual rating” task; moving bar replicated for each subject the specific pain rating task temporal pattern). The dataset includes one GLM beta map per subject per stimulus type. The dataset was previously described in ^44^.

#### Dataset 2

Fifty-one (51) healthy, right-handed, adult subjects (age = 24 ± 2 years, 34 females). Subjects had no history of brain injuries, pain disorders, or psychiatric or neurological diseases. fMRI data was collected while subjects received painful heat stimuli on the right foot dorsum using a CO_2_ laser, as well as tactile stimuli to the same area using electrical stimulation. Stimuli were not delivered at the same time. Perceived intensities were recorded for every stimulus and only the stimuli with matched perceived intensity for painful heat and touch were selected for GLM analysis. The dataset includes one activation map per subject per stimulus modality – painful heat and touch. The dataset was previously described in ^5,45^.

#### Dataset 3

Fourteen (14) healthy, right-handed, adult subjects (age = 20–36 years old, 6 females). fMRI data were collected while subjects received painful heat stimuli on the right foot dorsum using a CO_2_ laser, tactile stimuli to the same area using electrical stimulation, visual stimuli using a white disk presented above the right foot, and auditory stimuli delivered via pneumatic earphones. Stimuli were not delivered at the same time. Perceived intensities were recorded for every stimulus and only the stimuli with matched perceived intensity across the four modalities were selected for GLM analysis. The dataset includes one activation map per subject per stimulus modality – painful heat, tactile, auditory, and visual. The dataset was previously described and published in ^5^.

#### Dataset 4

Thirty-three (33) healthy, right-handed, adult subjects (age = 28 ± 9 years, 22 females). Subjects had no history of pain, psychiatric, or neurological disorders. fMRI data was collected while subjects received thermal stimuli that varied in one-degree Celsius increments across six temperatures from 44.3 up to 49.3°C. Subjects then evaluated each stimulus as warm, and scored it from 0, not perceived up to 99, about to become painful, or as painfully hot, and scored it from 100 (no pain) to 200 (worst imaginable pain). The dataset includes an average GLM activation map per subject per stimulus temperature, as well as the corresponding average stimulus ratings. When this dataset was applied dichotomously (pain vs. no pain), we averaged the brain activity maps from the painful and nonpainful conditions; we omitted subjects who had fewer than two brain activity maps for each condition, resulting in 29 subjects for dichotomous ratings. The dataset was previously described in ^4,46^.

#### Dataset 5

Two-hundred thirteen (213) healthy, adult subjects (age = 24 ± 7 years, 101 females). Subjects had no history of physical or mental health conditions. fMRI data were collected while subjects performed a voice localizer task. Forty blocks of vocal sounds (20) and non-vocal sounds (20) interspersed with periods of silence were presented while the subjects lay silent and passively listened with their eyes closed in the scanner. This dataset was previously described in ^47^. Raw fMRI data were downloaded from OpenNeuro (ds000158). We used minimal pre-processing for this study which was performed using the FMRIB 5.0.8 software library (FSL) ^48^, MATLAB2018a and in-house scripts. The following steps were performed: motion correction, intensity normalization, nuisance regression of 6 motion vectors, signal-averaged overall voxels of the eroded white matter and ventricle region, and global signal of the whole brain, and band-pass filtering (0.008–0.1 Hz) by applying a 4^th^-order Butterworth filter. All pre-processed fMRI data were registered to the 2×2×2 mm MNI152 template using a two-step procedure: the mean of preprocessed fMRI data was registered with a 7-degrees-of-freedom affine transformation (*x*, *y*, *z*, *α*, *β*, *γ*, and scale factor *k*) to its corresponding T_1_ brain (FLIRT); next, transformation parameters were computed by nonlinearly registering individual T_1_ brains to the MNI152 template (FNIRT). Combining the two transformations yielded a mapping from the preprocessed fMRI data to standard space. Task-related activation maps (vocal vs. silence, and non-vocal vs. silence) were derived from a whole-brain GLM regression analysis using the FMRIB Software Library (FSL) ^48–50^.

#### Dataset 6

Seventeen (17) healthy, adult subjects (4 women; age = 24 ± 5 years) partook in this study, which involved listening to a natural stimulus (5 min plot-driven audio story) and resting-state (first 5 min of 8 min scan) while under different levels of anesthesia ^7,8^. Sedation levels under propofol were determined by the Ramsey scale (awake, no propofol; light sedation, Ramsey = 3; deep sedation, Ramsey = 5; recover, Ramsey = 2, approximately 11 minutes after cessation of propofol) ^7^. We used minimal pre-processing for this study which was performed using the FMRIB 5.0.8 software library (FSL) ^48^, MATLAB2018a and in-house scripts. The following steps were performed: motion correction, intensity normalization, nuisance regression of 6 motion vectors, signal-averaged overall voxels of the eroded white matter and ventricle region, and global signal of the whole brain, and band-pass filtering (0.008–0.1 Hz) by applying a 4^th^-order Butterworth filter. All pre-processed fMRI data were registered to the 2×2×2 mm MNI152 template using a two-step procedure: the mean of preprocessed fMRI data was registered with a 7-degrees-of-freedom affine transformation (*x*, *y*, *z*, *α*, *β*, *γ*, and scale factor *k*) to its corresponding T_1_ brain (FLIRT); next, transformation parameters were computed by nonlinearly registering individual T_1_ brains to the MNI152 template (FNIRT). Combining the two transformations yielded a mapping from the preprocessed fMRI data to standard space. Task-related activation maps (vocal vs. silence, and non-vocal vs. silence) were derived from a whole-brain GLM regression analysis using R.

### Decoder Construction and Evaluation

Brain activity maps were masked to include only neocortical grey matter voxels using the Harvard-Oxford neocortical mask thresholded at 0.5. For each contrast, we performed a voxel-wise paired *t*-test using two brain activity maps from each subject, resulting in a *t*-statistic for each voxel in the grey matter. The *t*-statistic map was then binned into deciles by |*t*|—decile 10 contained the highest absolute value *t*-statistics (the “most significant”) and decile 1 contained the lowest absolute value *t*-statistics (the “least significant”). These deciled *t*-statistic maps served as our decoders.

We evaluated the decoders (**D** ∈ ℝ^*p*×10^) by multiplying them with the brain activity maps of interest (**B**_***I***_ ∈ ℝ^*n*×*p*^) and no interest (**B**_***NI***_ ∈ ℝ^*n*×*p*^), for *p* voxels and *n* subjects. This resulted in two matrices of dot products between the decoders and brain activity maps: one matrix of dot products from the activity maps of interest (**R**_***I***_ = **B**_***I***_ **D**) and one matrix of dot products from the activity maps of no interest (**R**_*N****I***_ = **B**_*N****I***_ **D**). The columns of **R**_***I***_ and **R**_*N****I***_ were then compared to calculate an AUC via the Mann-Whitney U-statistic (*AUC* = *U*_1_⁄*n*^2^). That is, column 1 in **R**_*I*_ was compared with column 1 in **R**_*N****I***_, column 2 in **R**_*I*_ was compared with column 2 in **R**_*N****I***_, and so on for all 10 columns, producing 10 AUCs—one for each decile. In doing so, we treated the subjects as dependent for decoder training (paired *t*-test) but independent for testing.

We constructed and tested all decoders using the 0.632+ bootstrap method with 100 replicates, which provides unbiased estimates of out-of-sample performance ^51^. Briefly, the 0.632+ bootstrap was performed as follows:

1. Train and test a model using the original sample. Let the resulting AUC be called the “apparent” AUC, 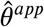.
2. Generate *b* bootstrap samples by resampling the original sample with replacement. Note, each bootstrap sample contains approximately 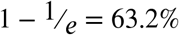 of the original sample. For each of the *b* bootstrap samples, we train the model on the bootstrap sample and test the model on the ~36.8% of individuals not part of the bootstrap sample. Let this AUC estimate be the “leave-one-out” (out-of-sample) bootstrap AUC, 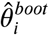.
3. Average the resulting out-of-sample bootstraps, 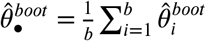.
4. Obtain the 0.632+ estimate.
  a. Calculate the relative overfitting rate,

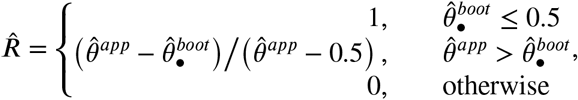

where 0.5 indicates no information in the decoder.
  b. Calculate the weight to adjust the 0.632 estimate,

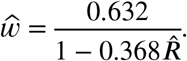
  c. Calculate the 0.632+ estimate,

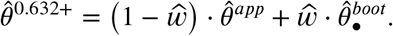

Note, we did not adjust 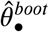 with 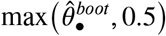 in 4c as is common ^51,52^, since this would create a floor effect such that 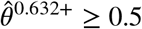, which would downwardly bias our variance estimates in the next step. This results in 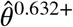 estimates that are identical to estimates with the adjustment when 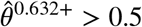, but allows our estimates to dip below chance (AUC = 0.5) since it removes the floor effect.

Variances and covariances of the AUCs were estimated using a nested bootstrap with 500 replicates in the outer loop and 100 replicates in the inner loop ^52^, totaling 500×100 = 50,000 replicates of each study. All inner and outer bootstraps were performed on the subject level. This sampling was carried out on Northwestern University’s High Performance Computing clusters (Quest), which took ~12 hours to complete using 50 cores.

### Meta-analysis

We performed a single-paper meta-analysis to consolidate our results ^53^. First, all AUCs were “squeezed” or shrunken toward 0.5 to avoid boundary effects ^54^,

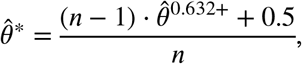

where *n* was the total number of brains used (i.e., twice the number of participants). Next, the 0.632+ bootstrap-estimated AUCs and their bootstrapped replicates were logit-transformed, and the logit-transformed bootstrap replicates were used to generate a 100×100 variance-covariance matrix of sampling errors. The logit-transformed AUCs were used as the response variable in a multivariate, multilevel linear meta-regression ^55^. This allowed for within-study dependence to be properly accounted for, including the dependence between deciles in a single contrast (e.g., decile 1 and decile 2 in Study 1) and the dependence between contrasts in Study 3 (e.g., decile 1 in touch vs. pain and decile 2 in visuomotor vs. pain). We were principally interested in the effect of decile on discrimination performance; we treated decile continuously and used it as a linear moderator (fixed-effect). Similarly, decile was treated continuously in the random-effect specification, wherein contrasts were nested within studies.

### Across-Subject Decoding with Principal Components Analysis (PCA)

To analyze the across-subject variance-covariance structure’s task-relevance, we performed PCA on the pain beta maps from Study 2 using singular value decomposition on a column-wise *z*-scored **B**_***I***_. To limit the number of principal components (PC) in our decoding analysis, we performed parallel analysis by generating surrogate data—100 null datasets—and calculating the variance explained by each null component, against which we compared our observed explained variance. Within each of the surrogate datasets, we performed discrete Fourier transforms on each beta map, scrambled their phases, and performed the inverse Fourier transform. This enabled us to maintain identical spatial frequency content and similar autocorrelation functions to the original beta maps. We used the number of real PCs that fell above the noise floor as determined by the parallel analysis.

After establishing with the whole sample that 4 PCs fell above the noise floor, we used the bootstrap 0.632+ to fit PC-based decoders. In each training set, we column-wise *z*-scored 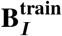 and performed PCA to obtain the top 4 PCs. We then independently *z*-scored the brain activity maps in the test set, 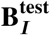 and 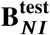, projected them into the four-dimensional PC space of 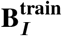, and calculated their respective Kullback-Liebler divergence from 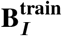 :

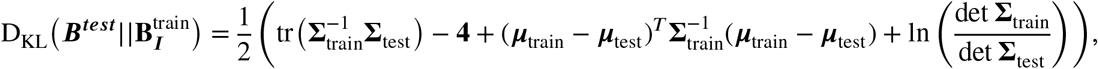

where ***μ*** and **Σ** were calculated in the PC space. A geometric depiction of this operation can be seen in Fig 3. If 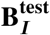 had a lower KL than 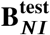, that test sample received a 1; otherwise, the test sample received a 0. The final bootstrap 0.632+ estimate thus represents the expected out-of-sample classification accuracy, not AUC.

### Perturbations

#### Noise

Since voxels with low signal-to-noise ratios (i.e., low *t*-statistics) were capable of decoding, we aimed to evaluate this finding’s boundary conditions. Each brain activity map contains a correlation coefficient *r_i_* for each voxel *i*, along with a *t*-statistic *t_i_*. We started with a brain of *t*-statistics, to which we added Gaussian noise 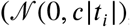, where *c* ∈ {0,1,2,3,4,5}). This procedure ensured that the noise added to each voxel was proportional to its signal-to-noise ratio to avoid biasing the regions with high signal. The *t*-statistics with added noise were then converted to Pearson’s *r*, on which we performed the decoding. Noise was added within each 0.632+ bootstrap replicate such that the resulting AUCs were averaged over 500 iterations (as opposed to 100 for other analyses) of added noise.

#### Voxel Sampling

Since the ability to decode with low signal-to-noise voxels likely arises from integrating over so many small bits of information, we randomly subsampled voxels. The full decoders contained approximately 10,735 voxels per decile, which could come from anywhere within our grey matter mask. We built and assessed decoders by subsampling the brain activity maps, such that the resulting decile-based decoders contained 100, 250, 500, 1000, 2500, 5000, 7500, and 10735 voxels each. Voxels were sampled within each 0.632+ bootstrap replicate such that the resulting AUCs were averaged over 500 iterations (as opposed to 100 for other analyses) of sampled voxels.

### Anatomical Specificity

#### Neocortex, Subcortex, and Cerebellum

Neocortical, subcortical (thalamus, striatum, hippocampus, and amygdala), and cerebellar grey matter voxels were extracted from each brain activity map. The neocortical grey matter mask contained 112,651 voxels; the subcortical mask contained 6,882 voxels; and the cerebellar cortex mask contained 17,142 voxels. Since decoding power is sensitive to the number of voxels, we randomly subsampled 6,882 voxels (or fewer for studies that were further masked) from each mask to control for voxel number. This subsampling was completed within each 0.632+ inner bootstrap replicate.

#### Neocortical Grey Matter, White Matter, and Cerebral Spinal Fluid

Neocortical grey matter (GM), white matter (WM), and cerebral spinal fluid (CSF) voxels were masked using the Harvard-Oxford atlas with conservative thresholds: 112,651 for GM; 61,324 for WM; and 1,926 for CSF. Within each study, we controlled for the number of voxels by resampling 1,925 voxels (since 1,926 < 61,324 < 112,651) from GM and WM within each bootstrap run.

### Anesthesia Decoders

The anesthesia dataset employed a naturalistic audio stimulus and thus does not have a task vector associated with it. Moreover, this was the only task performs. As such, we compared each anesthesia level’s task (naturalistic listening) to resting state. To facilitate this, we used the average auditory cortex activity from the training sample as the task vector. Of course, this analysis is circular within the training sample, but because the training sample’s brain activity was used as the vector in the testing sample and decoding was assessed based on the resulting brain activity maps, the testing is not circular. To extract the auditory cortex vector, we defined a region of interest (ROI) based on the Neurosynth association map for “auditory”, which was thresholded using a *z*-score of 12.

#### Decile Decoders

Decile-based decoders for the anesthesia dataset were created in a similar manner to the other datasets. To summarize the performance within each decile, we fit a single generalized least squares model on the logit-transformed AUCs from all anesthesia states, **y**. To do so, all AUCs were “squeezed” towards 0.5 like they were for the meta-analysis. Our weight matrix, **W**, was defined as the inverse of the variance-covariance matrix of the logit-transformed bootstrapped AUCs, 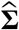. The parameter estimates, 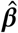, and their standard ewrerorerscalculated as

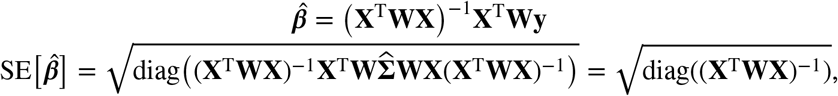

where **X** is the design matrix,

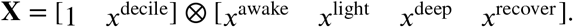

This differs from standard weighted least squares (a diagonal weight matrix) but has more favorable properties since it accounts for covariation.

#### Region of Interest Decoders

We created posterior cortex, anterior cortex, visual cortex, and motor cortex ROIs using the Harvard-Oxford neocortical atlas, thresholded at 25%. The posterior cortex was specified as areas 20–22; the anterior cortex, areas 1, 4, and 5; the visual cortex, areas 36, 40, and 48; and the motor cortex, area 7. In addition, we used the same auditory cortex ROI as described above. Again, the averaged auditory ROI time course from the training sample was used as the task vector. A decoder was created using the *t*-statistics from each ROI (without deciles), which was fit and tested using the same approach as the decile decoders.

#### Functional Connectivity

Pearson correlation coefficients were used to calculate within- and inter-subject (ISFC) functional connectivity between the auditory cortex and the other ROIs, during both the auditory task and rest within each level of anesthesia. Within-subject functional connectivity was calculated by averaging the time course between all voxels within each ROI, calculating the correlation between the auditory ROI and all other ROIs, converting from Pearson’s *r* to Fisher’s *z*, averaging across participants, and then converting back to Pearson’s *r*. ISFC was calculated similarly to previous work ^56^. When calculating subject 1’s ISFC, we correlated subject 1’s auditory cortex time course with the average time course of, for example, posterior cortex from subjects 2–*n*. This was repeated for all subjects and the resulting auditory cortex-posterior cortex ISFCs were averaged using Fisher’s *z* to obtain the final estimate of the auditory cortex-posterior cortex ISFC. Again, this was repeated for the anterior, visual, and motor cortex; ISFC was also measured between auditory cortices across all subjects.

## Footnotes

† By *perception*, we refer to the conscious experience resulting from sensory input. This experience is the product of and thus includes memory, attention, and expectations.

‡ Here, we use *information* not as an inference regarding neuronal function, but rather from a decoding perspective relating to the nature and specificity of the variance that can be extracted from our proxies of neuronal activity.

## Acknowledgments

We would like to thank Drs. Christof Koch, Todd Parrish, and Lucas Pinto, in addition to the Apkarian Lab members, for their thoughtful comments and feedback. Our acknowledgment does not imply endorsement of our views by these colleagues, and we remain solely responsible for the views expressed herein.

