## Supplement for "Widespread, perception-related information in the human brain scales with levels of consciousness"

Vigotsky *et al.* (2023)

**Table S1.** ***Studies and contrasts analyzed.***

| **Study** | ***n*** | **Contrasts** |
|---|---|---|
| Baliki *et al*. (*1*) | 15 | Pain vs. visuomotor |
| Liang *et al*. (*2*) | 51 | Pain vs. touch |
| Liang *et al*. (*2*) | 14 | Pain vs. touch |
| | | Pain vs. audition |
| | | Pain vs. vision |
| | | Touch vs. audition |
| | | Touch vs. vision |
| | | Audition vs. vision |
| Wager *et al*.(*3*) | 31 | Pain vs. heat |
| Pernet *et al.(4)* | 213 | Vocal vs. non-vocal sounds |
| Naci *et al.* (*5*) & Kandeepan *et al*. (*6*) | 17 | Awake: audition vs. rest |
| | | Light sedation: audition vs. rest |
| | | Deep sedation: audition vs. rest |
| | | Recovering: audition vs. rest |

The first 5 studies were used to generate task contrasts, which assessed the extent to which individual voxels were preferentially correlated with one of the two tasks. Univariately, these contrasts indicate the presence of information in the cortex, subcortex, and cerebellum via the univariate signal-to-noise (*t*-statistic), which we report in deciles. The first study (Baliki et al.) was also used to compare task-related information content between cortical grey matter, white matter, and ventricles. The last study (Naci et al., Kandeepan et al.) was used to examine the dependence of brain information content on level of sedation, both as a function of univariate signal-to-noise level and across five broad brain regions.

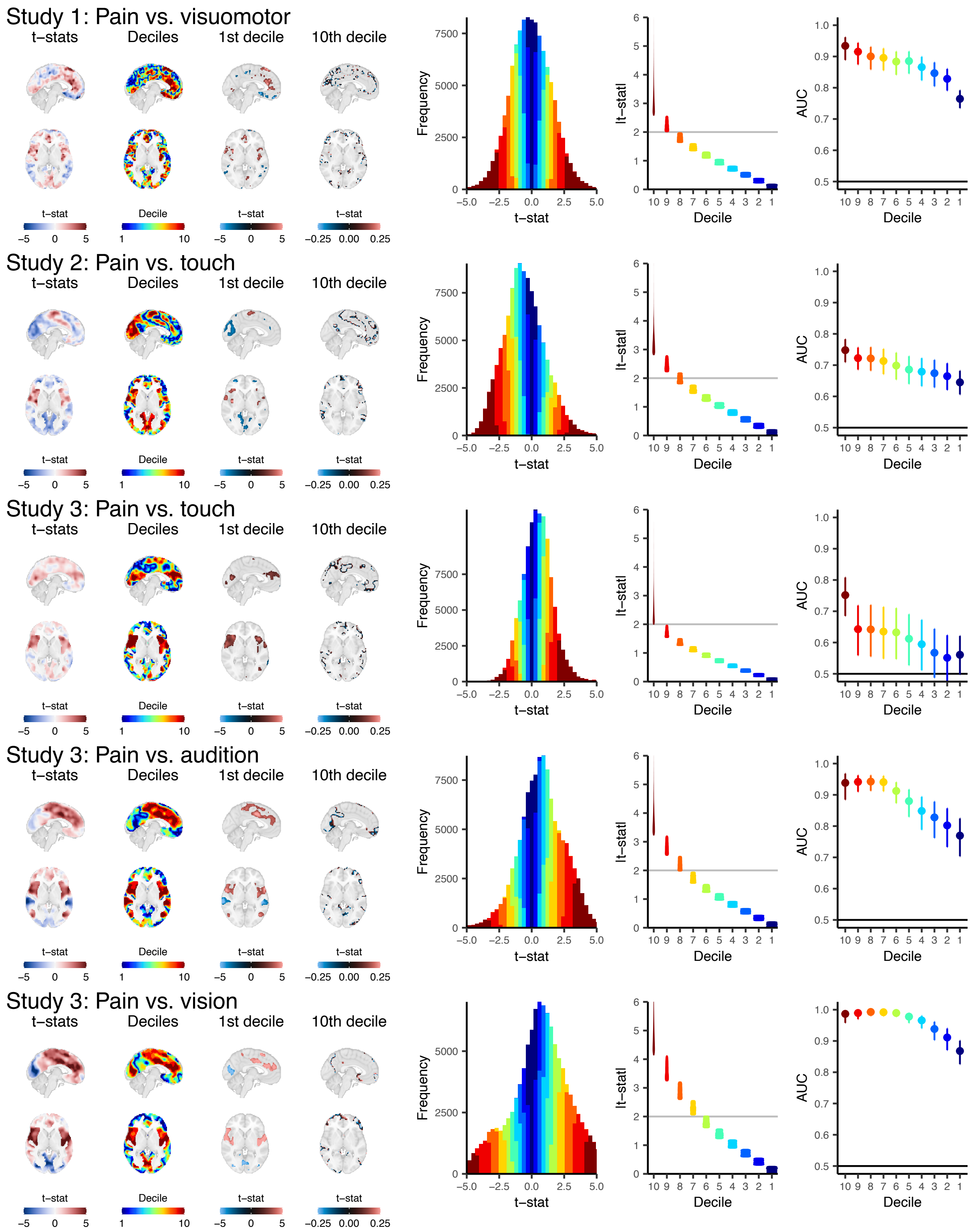

Study 1: Pain vs. visuomotor
Study 2: Pain vs. touch
Study 3: Pain vs. touch
Study 3: Pain vs. audition
Study 3: Pain vs. vision
t–stats
Deciles
1st decile
10th decile
t–stat
Decile
−5 0 5
1 10
−0.25 0.00 0.25
Frequency
t–stat
|t–stat|
Decile
AUC

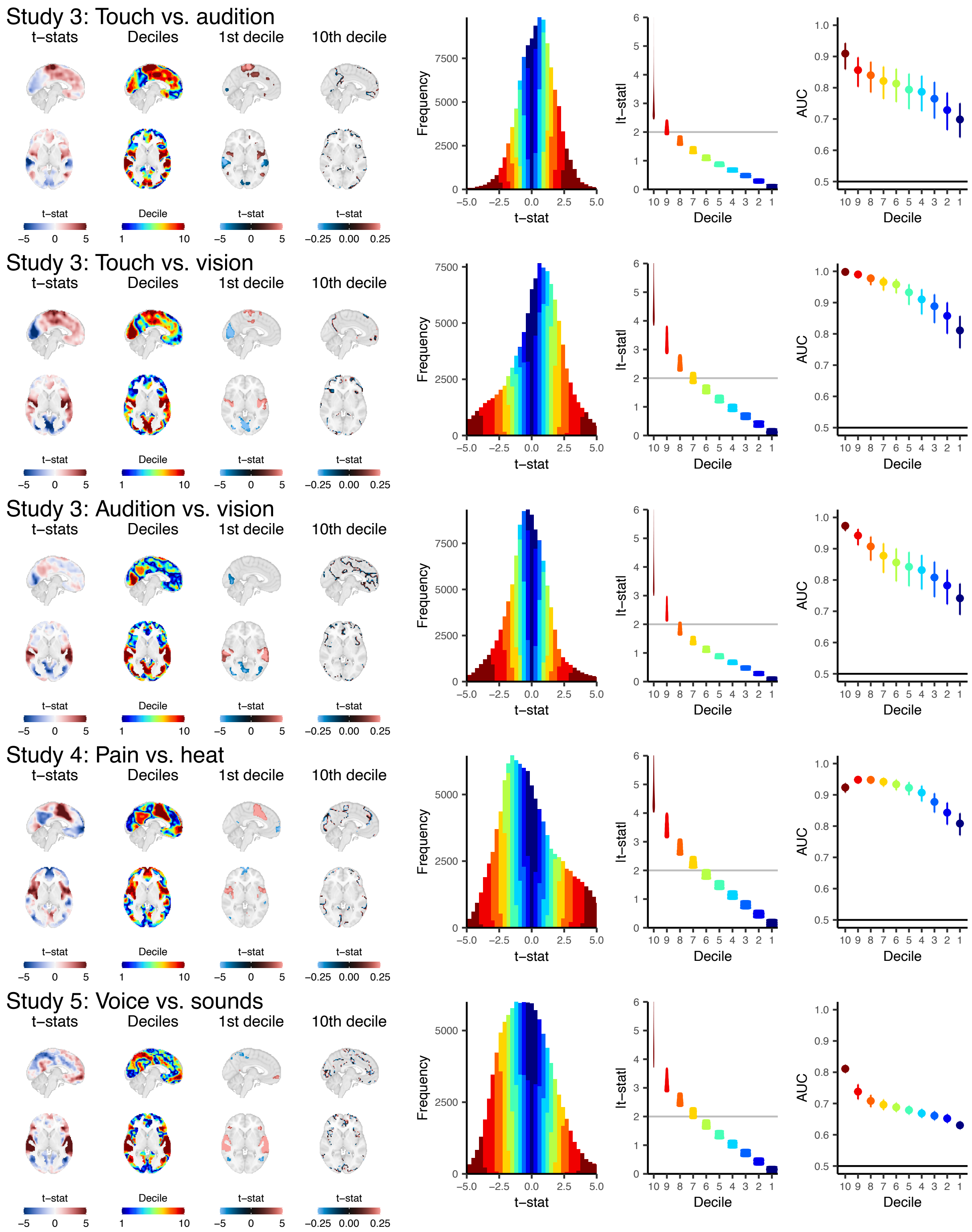

**Figure S1.** ***Contrast maps (t-statistics) and their associated deciles and discrimination performance.***
Voxel-wise contrasts were performed using the brain activity maps (beta maps) from each pair of tasks within each study. For each contrast, we have the raw *t*-statistic map (leftmost), its associated

deciles (second from the left), the *t*-statistics in the highest decile (second from the right), and the *t*-statistics in the lowest decile (rightmost). Also depicted is the histogram of *t*-statistics, colored by decile, in addition to their binning and respective discrimination performance (AUC). Note that the *t*-statistics maps have spatial autocorrelation and thus the deciles have high amounts of contiguity. This results in the highest deciles tending to show up in clusters, while the lowest deciles tend to be where the *t*-statistics are transitioning in sign (e.g., from positive to negative). The grey line in the |*t*-stat| plot indicates $t \approx 2$, approximately the uncorrected minimum threshold for "statistical significance," which is used to identify regions that are reliably related to a task in univariate analyses. Error bars on AUCs indicate bootstrapped standard errors.

In our decoding analyses, all voxels within a decile were used to decode, with which a weighted sum was used to contrast the two tasks (using an unbiased, out-of-sample approach; see *Methods*). This simple, and arguably naïve, multivoxel decoding approach uncovered task-related information essentially everywhere in the neocortical grey matter. Across all 10 contrasts, derived from 5 distinct studies (data collected in different labs, $N = 293$ subjects), task discrimination (AUC) was consistently above chance across task pairs and deciles, except in case where CIs were compatible with chance (Study 3, Pain vs. touch).

At the highest decile, as illustrated in two brain slices ($10^{th}$ Decile), identified voxels are localized to contiguous blobs in brain regions commonly known to be involved in the contrasted tasks. For example, in Study 3's Audition vs. Vision contrast, the $10^{th}$ Decile identifies auditory and visual cortices as high positive (red) and negative (blue) *t*-value regions. Using all the *t*-values in the $10^{th}$ Decile as decoder weights, with which we obtain almost perfect discrimination, an AUC $\approx$ 1.0. For the same study and contrast—Study 3, Audition vs. Vision—the $1^{st}$ Decile identifies voxels scattered throughout the cortex. Despite the *t*-statistics being close to zero, the decoder still adequately discriminates between conditions, although less accurately, AUC $\approx$ 0.7. Note, the number of voxels comprising each decile (decoder) is always the same within each study, and ranges from 5,768 to 11,265, where the total number of cortical grey matter voxels ranges from 57,787 to 112,651, which is dictated by the conjunction of each study's common mask and neocortical grey matter voxels.

The univariate *t*-statistic distributions are unique for each study and each contrast pair; their uniqueness likely depends on many parameters that we are unable to control or explore. The 10 deciles cover all grey matter voxels in the neocortex. Yet, given the unique *t*-statistic distribution for each study and contrast, the spatial patterns for each decile are task- and contrast-specific, although their extent of clustering diminishes continuously down to individual voxels at decile 1. In all but one contrast, we observed presence of above chance information at all deciles, demonstrating the presence of task-specific information across all voxels of the neocortex that could only be captured by our decoders, which are weighted sums across 10% of the voxels of the neocortical grey matter. Like the *t*-statistic distributions, and in a related fashion, the peaks and slopes of the AUC-decile curves are also study- and contrast-dependent. This heterogeneity is quantified in our meta-analysis (Table S2). Despite this heterogeneity, our meta-analysis of discrimination performance across all the contrasts provides an overview of the decay of information content within the neocortex as a function of univariate signal-to-noise ratios (deciles of *t*-statistics; see Fig 1 and Table S2). The variance parameters indicate the presence of task-specific information throughout the cortical grey matter between the tasks and contrasts included in the meta-analysis.

**Table S2.** ***Meta-regression of discrimination performance across studies and contrasts.***

| **Fixed-effects** | **Estimate ± SE** | **$CI_{95\%}$** | ***z*-value** | ***P*-value** |
|---|---|---|---|---|
| Intercept | 0.9301 ± 0.1588 | (0.6189, 1.2413) | 5.8672 | < 0.0001 |
| Decile | 0.1325 ± 0.0326 | (0.0687, 0.1963) | 4.0692 | < 0.0001 |
| Moderators: QM(df = 1) = 16.5586, *P* < 0.0001 | | | | |
| **Random-effects** | **Variance estimate** | | | |
| ***Level:*** *Study* | | | | |
| Intercept | 0.0288 | | | |
| Decile | 0.0023 | | | |
| ***Level:*** *Contrast* | | | | |
| Intercept | 0.1434 | | | |
| Decile | 0.0043 | | | |
| Residual heterogeneity: QE(df = 98) = 772.0020, *P* < 0.0001 | | | | |

We performed a multivariate, mixed-effects meta-regression of decoding performance across deciles using the 10 contrasts shown in Fig S1. Deciles were shifted to start at zero instead of one (deciles 0–9 instead of 1–10) to improve interpretability, meaning that the intercept corresponds to the first decile. Deciles were modeled within contrasts, which were nested within studies. Since the model was fit on logit-transformed AUCs, the coefficients must be inverse logit-transformed. Thus, the intercept (decile 1) corresponds to an AUC of 0.72. AUCs are expected to increase by 0.13 logit units for each decile, rising to 0.9 by decile 10. Since one of the studies contained multiple contrasts, we could begin decomposing the random-effects into study- and contrast-level variances. In general, contrast-related variance seemed to dominate study-related variance, and much of this variance was accounted for by intercept as opposed to slope variability. However, these random effects should be interpreted with caution since they strongly rely on one study (Study 3). Nevertheless, the meta-regression enabled us to summarize our findings with a single, digestible model indicating the ability to decode across deciles from several studies and contrasts.

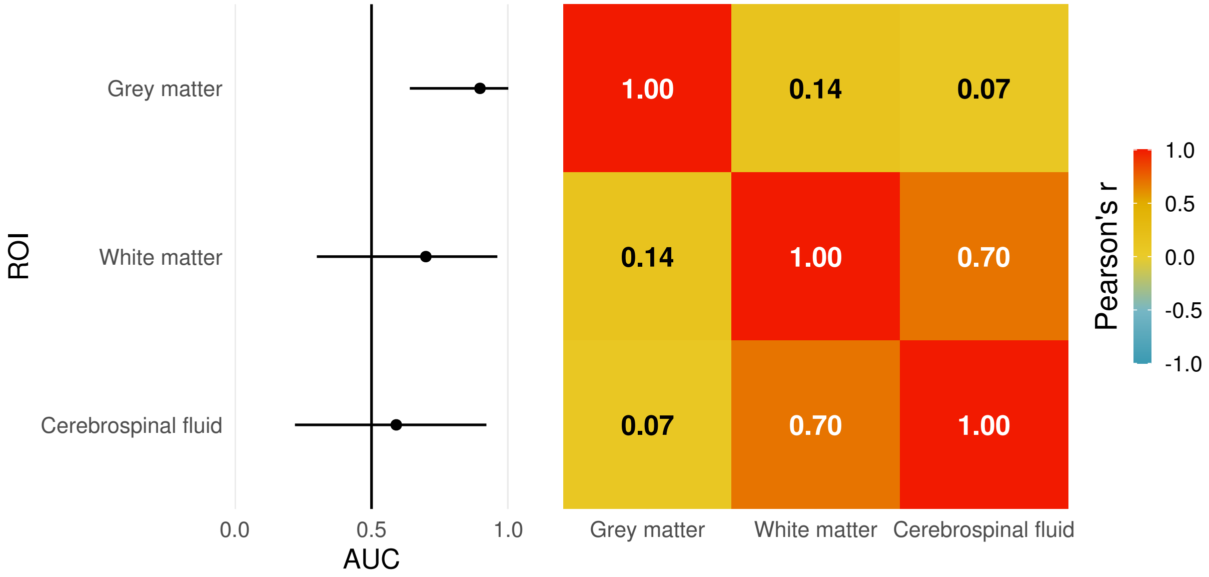

**Figure S2.** ***Signal in cortical grey matter (GM) is greater than that in white matter (WM) and cerebral spinal fluid (CSF), is minimally correlated with WM and CSF.***
Left, AUCs and their 95% CIs for GM, WM, and CSF in Study 1. GM has the greatest point estimate, followed by WM, then CSF. However, the CIs are wide, especially for WM and CSF; thus, we cannot rule out that these regions contain information.
Right, despite not being able to rule out that WM and CSF contain task-specific information, we assessed how their AUCs relate to GM's AUCs via the covariance (correlation) of the logit-transformed AUC bootstrap replicates. WM and CSF are minimally correlated with GM but are strongly correlated with one another, suggesting GM's information content likely has a different provenance than that of WM and CSF.

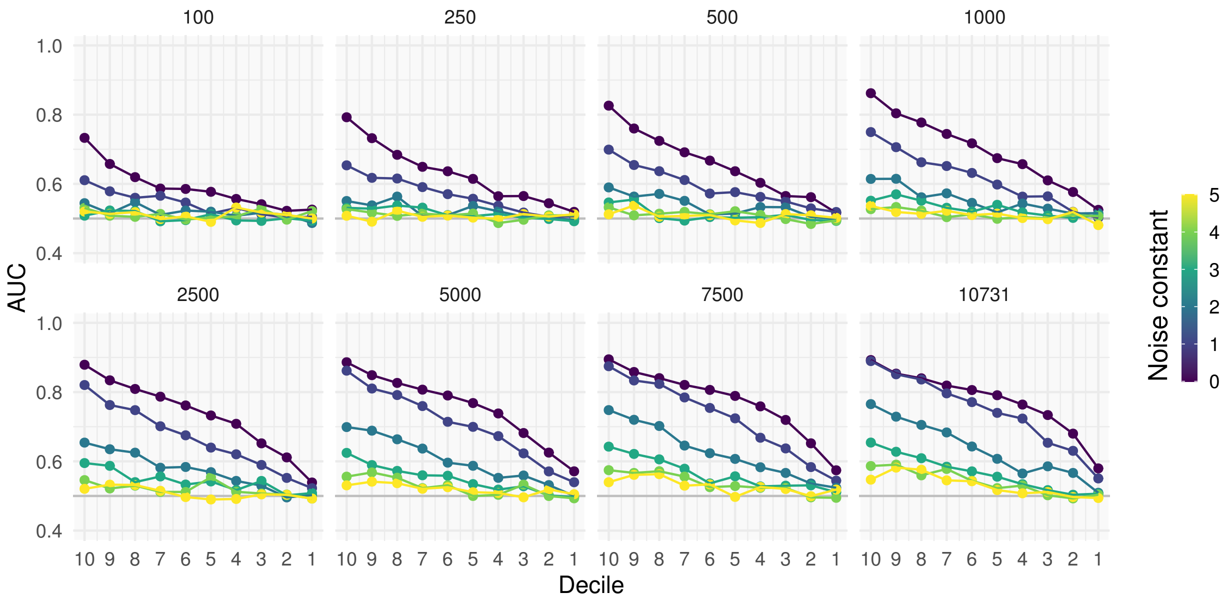

**Figure S3.** ***Signal is sensitive to the number of voxels and degrades when Gaussian noise is added to the data.***

In our primary analysis, each decoder contained 10% of the voxels in the neocortex. We hypothesized that their ability to decode well was at least in part due to the number of voxels contained in each decoder. In the extreme case, a decoder from the 10$^{th}$ decile using only one voxel should still be able to decode well, since that voxel alone can, by definition, discriminate between the conditions. However, a single voxel from the 1$^{st}$ decile likely cannot decode on its own since it is approximately orthogonal. In other words, the decoding ability of the first decile comes from the integration of many small bits of information. As such, decoding performance should be sensitive to the number of voxels (amount of information) that we use. To assess this hypothesis, we randomly sampled different numbers of voxels to assess the dependence on voxel counts. For this, we used linearly registered data from Study 1 since the nonlinearly registered data had more spatial autocorrelation.

Each panel contains a different number of voxels that were randomly sampled within each bootstrap replicate. The first panel contains 100 voxels, the second panel contains 250 voxels, and so on. The last panel contains 10,731 voxels, and so these decoders cover the entire neocortex without any random sampling. Across panels, there is a systematic decrease in the ability to decode: the more voxels used, the better discrimination. However, it seems the voxel-dependence is also dependent on the decile. The 1$^{st}$ decile drops off quickly, while the 10$^{th}$ decile seems more robust, with it only really seeing an appreciable decrease in performance with 500 voxels (vs. 10,731).

In addition, we perturbed each decoder by adding noise that was proportional to the *t*-statistic. This effect also had a decile and voxel number dependence. For instance, AUCs with a noise constant of 0 and 1 were almost identical for the 10$^{th}$ decile with 10,731 voxels, but there was an appreciable difference between the two when decoding with 100 voxels. On the other hand, the lower deciles seemed to be much more sensitive to this added noise.

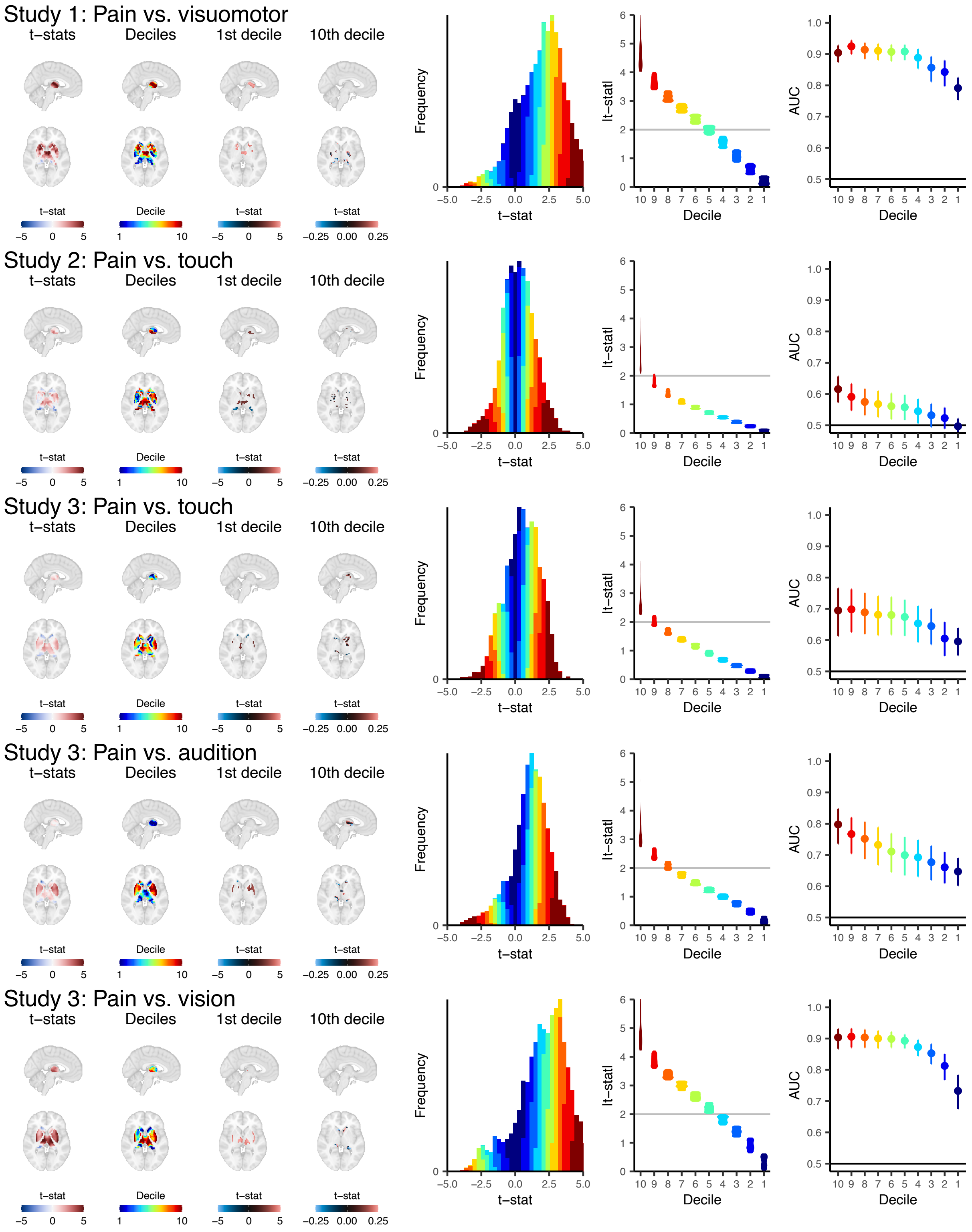
Study 1: Pain vs. visuomotor
t−stats
Deciles
1st decile
10th decile
t−stat
Decile
−5 0 5
1 10
−0.25 0.00 0.25
Frequency
t−stat
−5.0 −2.5 0.0 2.5 5.0
|t−stat|
Decile
10 9 8 7 6 5 4 3 2 1
AUC
1.0 0.9 0.8 0.7 0.6 0.5
Study 2: Pain vs. touch
Study 3: Pain vs. touch
Study 3: Pain vs. audition
Study 3: Pain vs. vision

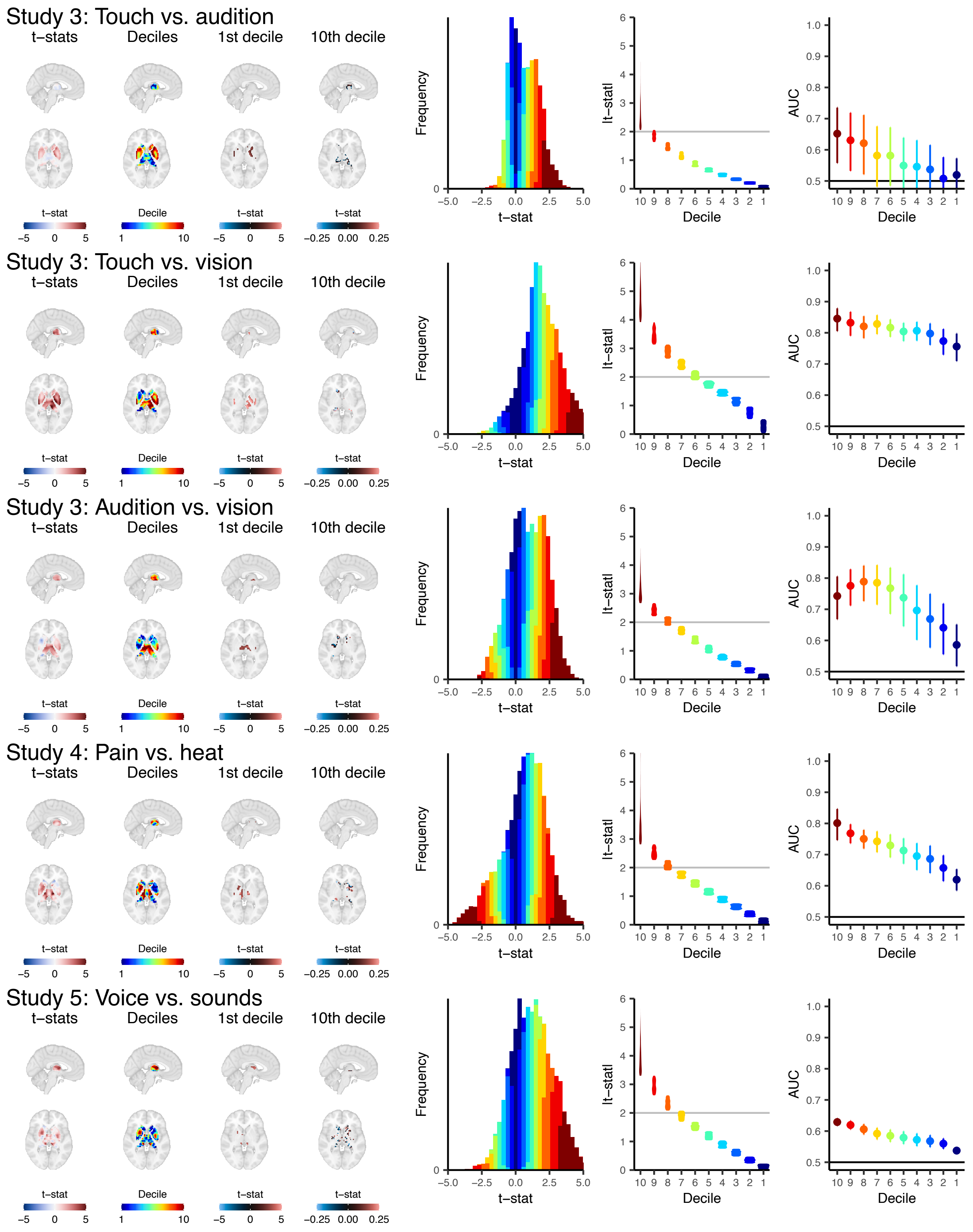

**Figure S4. *Subcortex contains signal across deciles.***
We built decile-based decoders for the subcortex in the same way we built them for the neocortex (see *Methods* and Figure S1). Our results for the subcortex are largely consistent with those of the

neocortex, except for one contrast (Study 3, Touch vs. audition), in that we were consistently able to recover task-specific information across nearly all deciles.

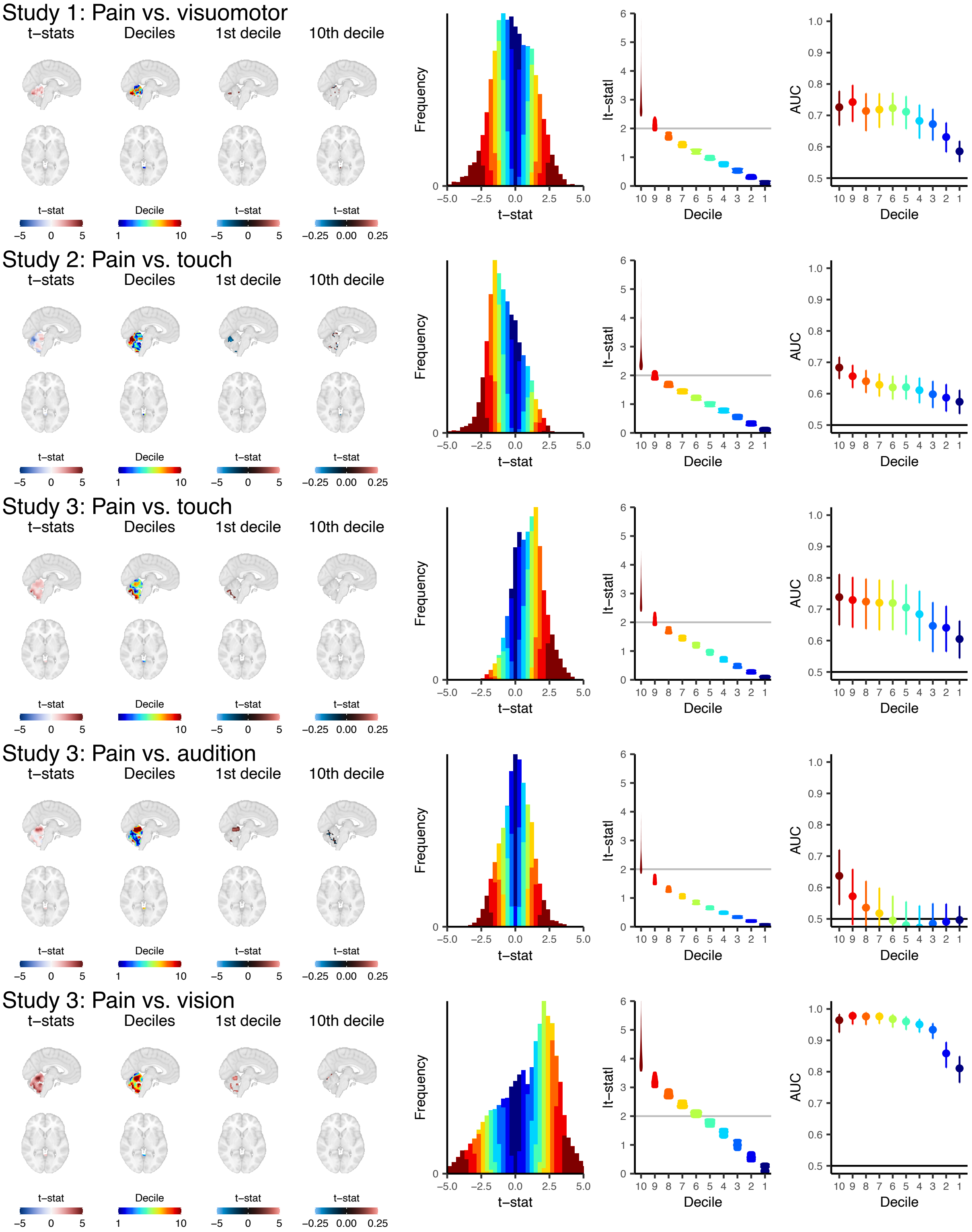

Study 1: Pain vs. visuomotor
Study 2: Pain vs. touch
Study 3: Pain vs. touch
Study 3: Pain vs. audition
Study 3: Pain vs. vision
t−stats
Deciles
1st decile
10th decile
t−stat
Decile
−5 0 5
1 10
−0.25 0.00 0.25
Frequency
t−stat
−5.0 −2.5 0.0 2.5 5.0
|t−stat|
Decile
10 9 8 7 6 5 4 3 2 1
AUC

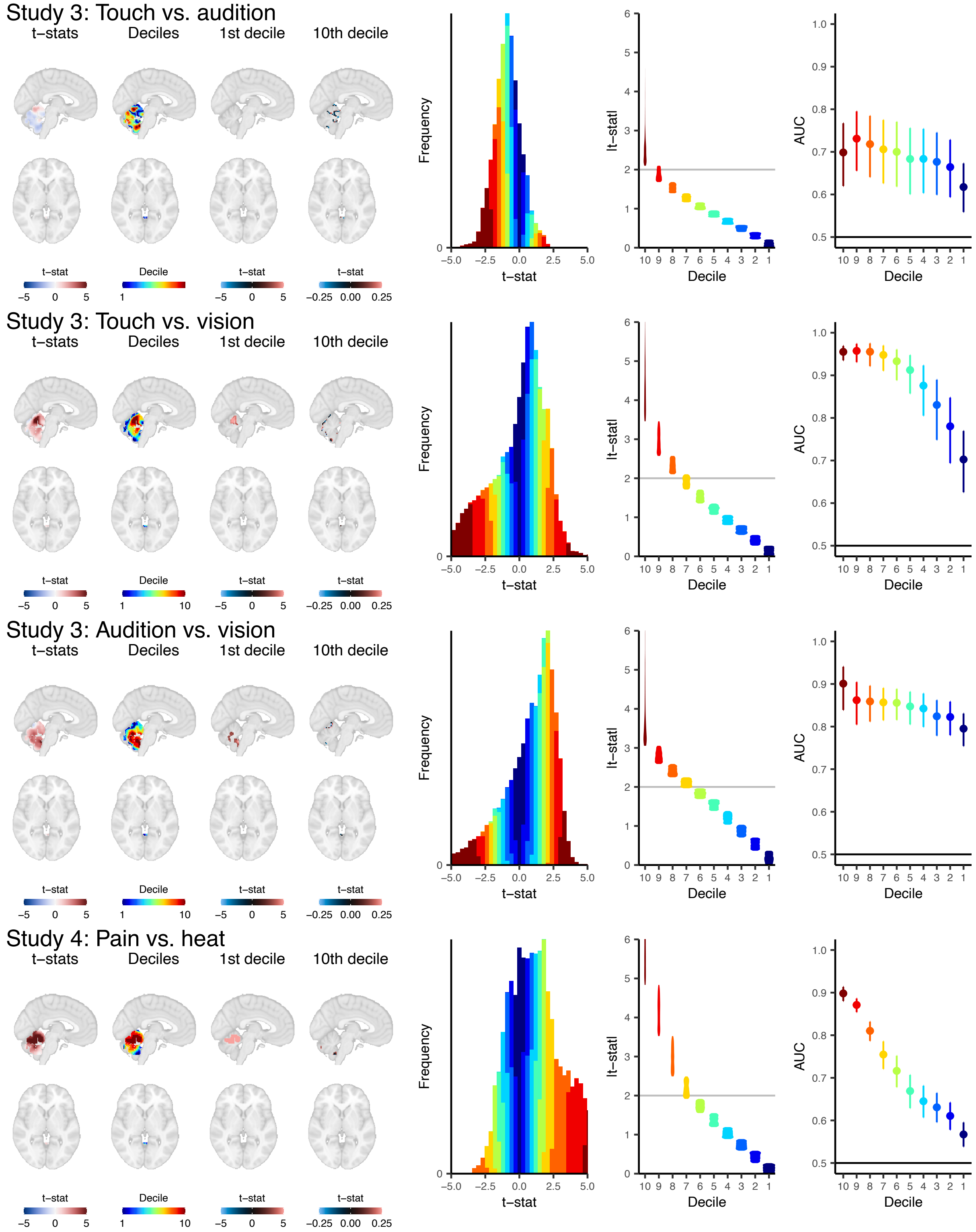

**Figure S5.** ***Cerebellum contains signal across deciles.***
We built decile-based decoders for the cerebellum in the same way we built them for the neocortex (see *Methods* and Figure S1). For this analysis, we discarded Study 5 since the cerebellum was cut off in most participants. Our results for the cerebellum are largely consistent with those of the

neocortex, except for one contrast (Study 3, Pain vs. audition), in that we were consistently able to recover task-specific information across nearly all deciles.

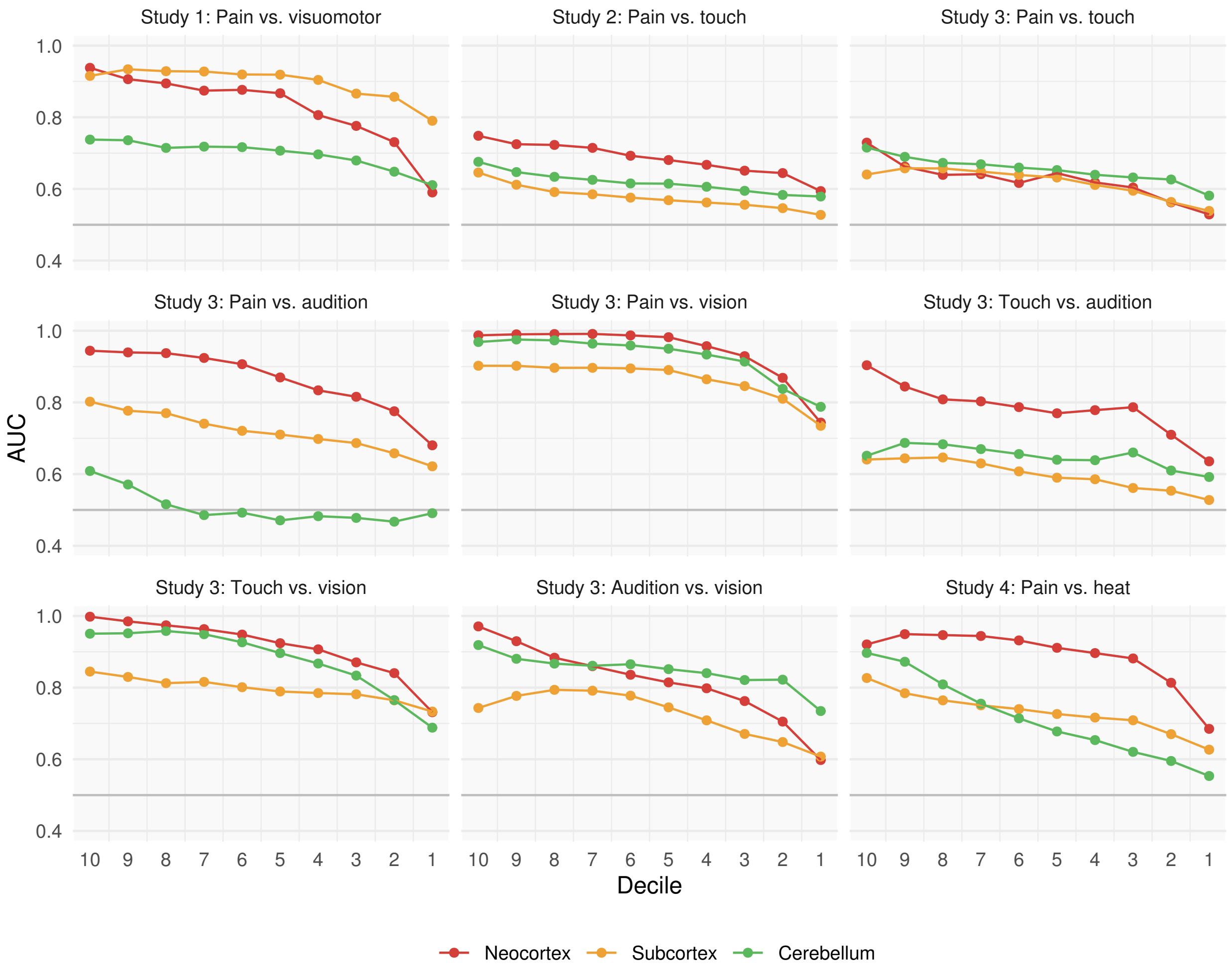

**Figure S6.** ***Voxel-matched neocortical, subcortical, and cerebellar decoding performance.*** Since decoding performance is dependent on the number of voxels used (Figure S3), we controlled for the number of voxels to compare the neocortex, subcortex, and cerebellum within each contrast. There was marked heterogeneity between studies. For instance, in Study 1, the neocortex and subcortex start at nearly the same point, but neocortex drops off more quickly. On the other hand, the cerebellum starts off with moderate performance and is steady across deciles, such that the neocortex has similar performance by the 1$^{st}$ decile. In contrast to Study 1, Study 4 shows a very different trend: the neocortex and cerebellum start off similarly, but the cerebellum drops off quickly and the subcortex decreases steadily. In Study 3, Pain vs. touch, all regions decode similarly across deciles, while in Study 3, Pain vs. audition, there are huge discrepancies between regions. Thus, there is marked heterogeneity in the patterns of information that our decoders captured for reasons that remain unclear.

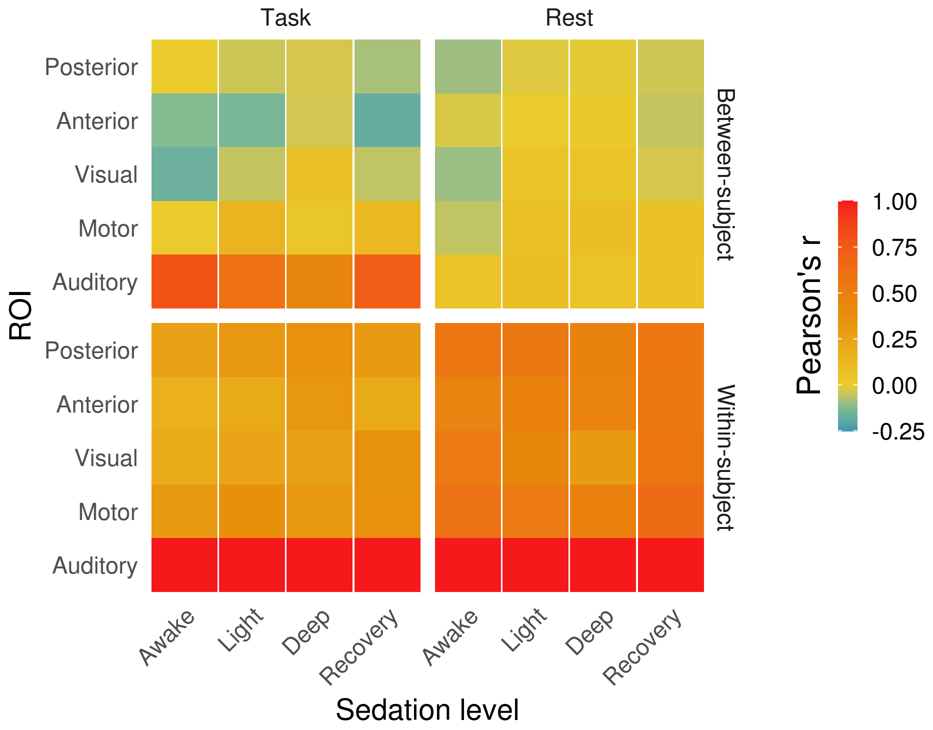

**Figure S7.** ***Within- and inter-subject functional connectivity with the auditory cortex across different sedation levels.***

We estimated two types of functional connectivity (FC) separately across different levels of sedation: within- and between-subjects. In both cases, the auditory cortex served as the "seed." Within-subject FC was calculated by, for example, correlating participant *i*'s auditory cortex with participant *i*'s posterior cortex. This was repeated for all *n* participants; the resulting correlation coefficients were converted to Fisher's *z*, averaged, and converted back to Pearson's *r*. In contrast, between-subject or inter-subject FC (ISFC) was calculated by, for example, correlating participant *i*'s auditory cortex with the average of all other participants' posterior cortex activity. Again, this was repeated for all *n* participants; the resulting correlation coefficients were converted to Fisher's *z*, averaged, and converted back to Pearson's *r*. As such, within-subject FC represents a very different construct than ISFC; ISFC presumably reflects *only* the task-dependent signal, while within-subject FC contains many other sources of covariance.

The original study from which these data came reported the effect of sedation and the task on within-subject FC (*5*), like our bottom panels, albeit with a different parcellation. Within-subject FC demonstrates relatively homogenous patterns, whereas ISFC varies much more appreciably as a function of both task and level of sedation. For the task, ISFC seems to "flatten" with greater levels of sedation and is relatively "flat" across conditions during rest. These results are consistent with our decoding performance: auditory cortex maintains task-specific information, but task-specific information degrades across the other cortices with sedation, in turn preventing our ability to decode.

Given these results and their constructs, it seems within-subject connectivity is less relevant than ISFC (*7*) when decoding across subjects.

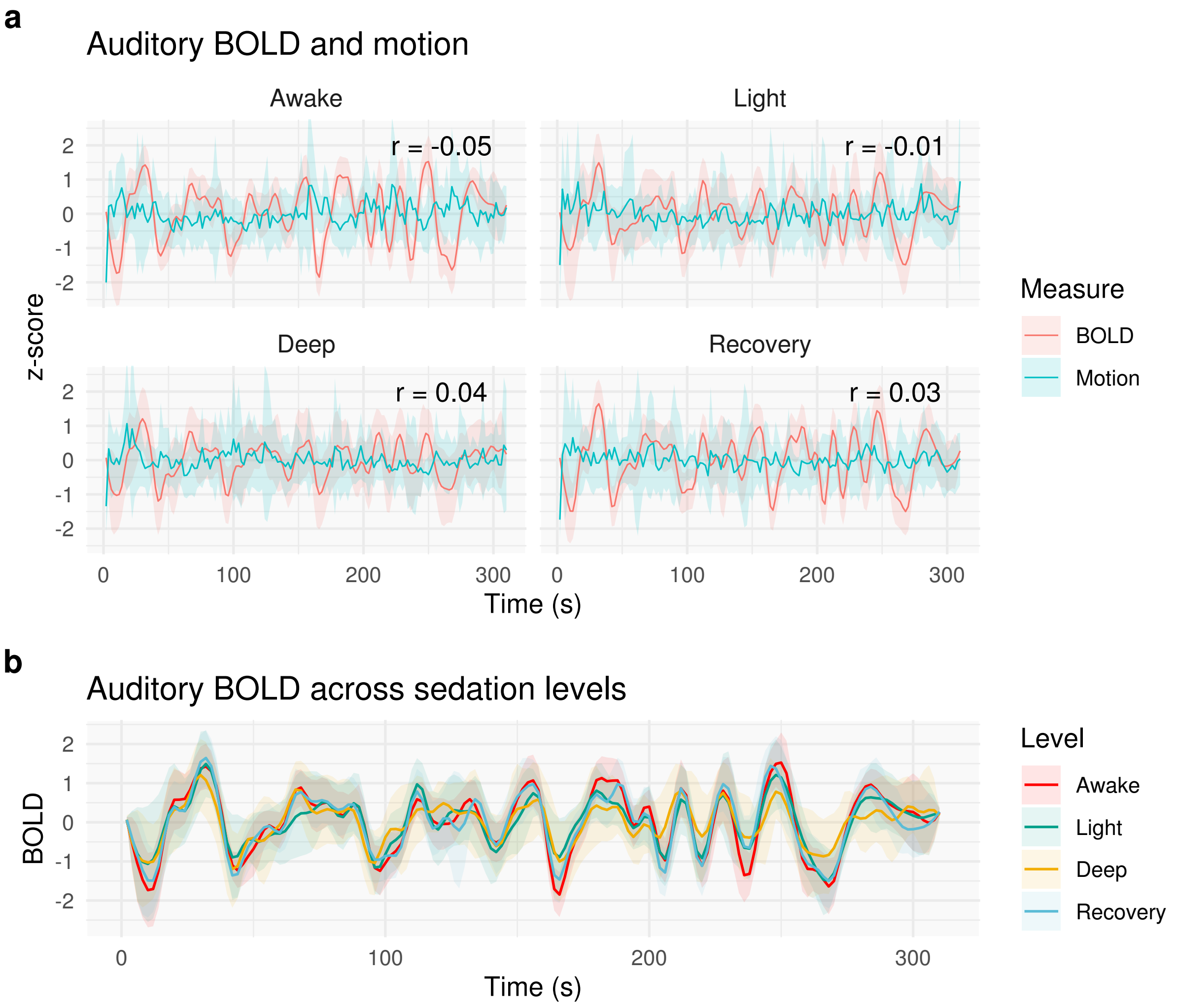

**Figure S8.** ***Auditory cortex BOLD activity and motion vectors have negligible temporal correlations in the sedation dataset, and auditory BOLD activity strongly correlates across sedation levels.***

**(a)** Depicted are $z$-scored vectors of the auditory cortex BOLD signal (red) and absolute frame displacement (blue). Even when participants are awake, the correlation between the BOLD signal and motion is negligible. Here, the correlations were calculated within-subject and averaged using Fisher's $r$-to-$z$ and then converted back to Pearson's $r$. **(b)** Average $z$-scored auditory cortex BOLD activity strongly correlates across sedation levels ($r$ = 0.85–0.96). Lines are the group mean and error ribbons indicate ± SD.

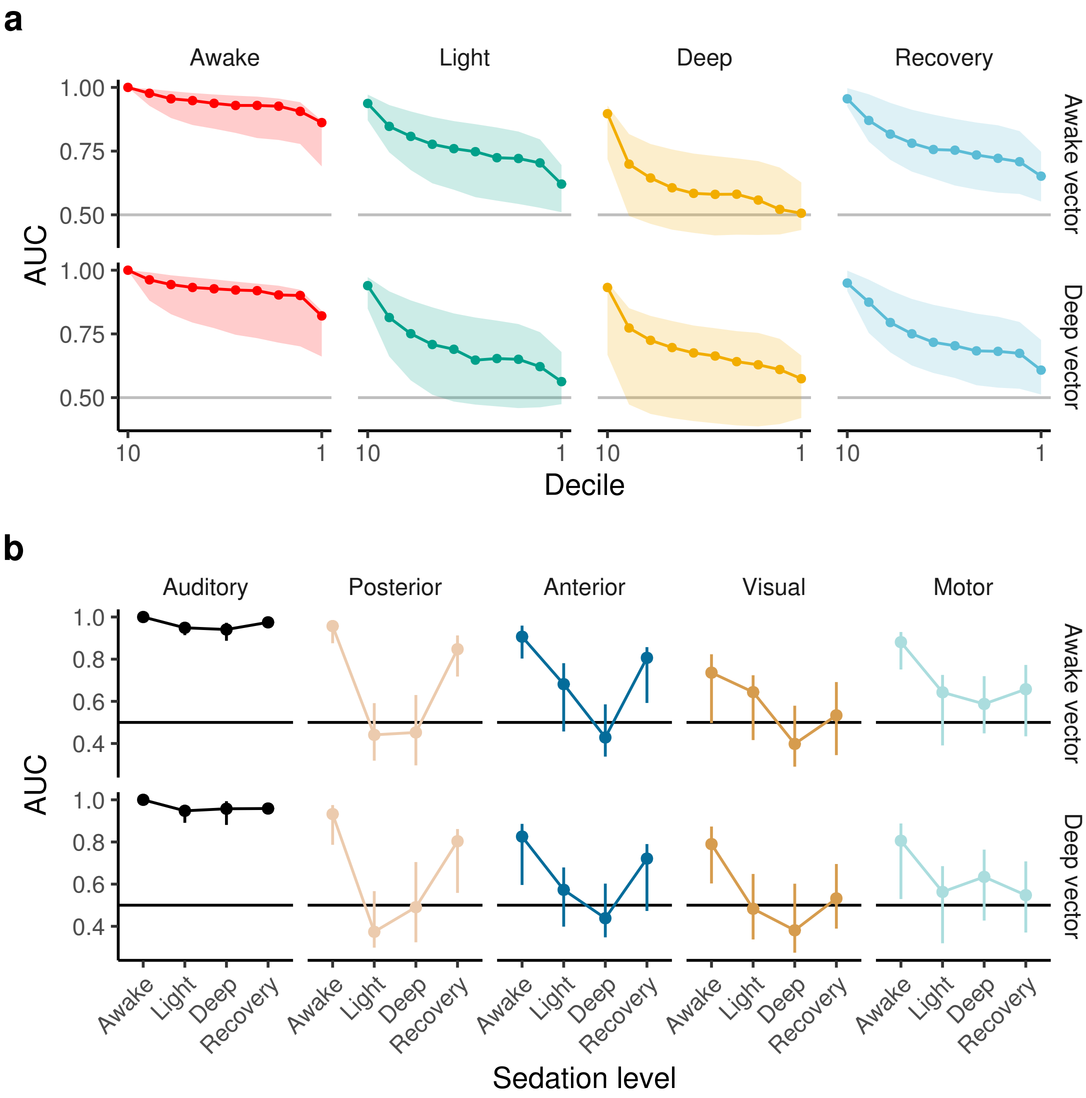

**Figure S9.** ***Results are similar when the task vector is derived from awake or deep auditory cortex activity.*** **(a)** Decile-based decoders perform similarly whether the task vector used is derived from auditory cortex activity when participants are awake (awake vector) or under deep sedation (deep vector). **(b)** ROI-based decoders perform similarly whether the awake vector or deep vector is used as the task vector. These results stress that our result is robust to task-related artifacts, such as stimulus salience, motion, etc. that can introduce shared variance across the brain. In particular, the deep sedation condition will have less task-related artifact since the participant is unconscious. This suggests our results are independent of that task-related artifact (e.g., motion), since the participant is not grossly reacting to external stimuli.

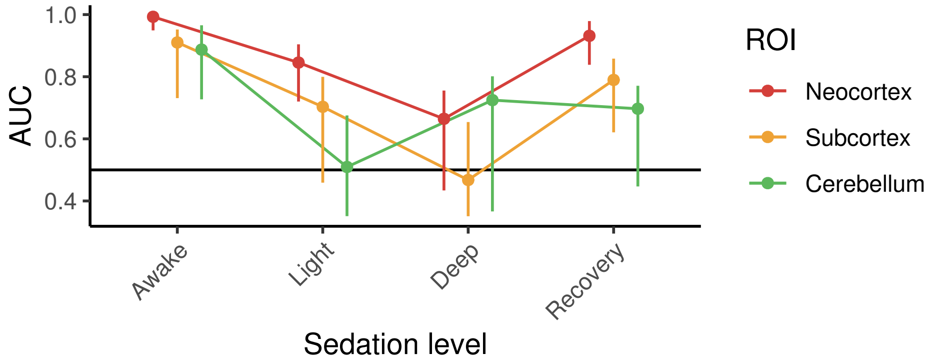

**Figure S10.** ***Neocortex, subcortex, and cerebellum across different levels of sedation, controlling for voxel counts.***
We decoded neocortex (minus auditory cortex), subcortex, and cerebellum across different levels of sedation. We controlled for number of voxels within each decoder, such that each instantiation contained 6,070 voxels. In general, these results followed a similar trend to our other analyses, with information tending to decrease with greater levels of sedation. This was even the case for the cerebellum, which has been posited to not be necessary for consciousness (*8*). Due to the small number of voxels and uncertainty associated with sampling these voxels, our CIs are wide, and thus, tracking the exact trend of each ROI is difficult ascertain.
